# Inversely proportional myelin growth due to altered *Pmp22* gene dosage identifies PI3K/Akt/mTOR signaling as a novel therapeutic target in HNPP

**DOI:** 10.1101/2021.11.08.467756

**Authors:** Doris Krauter, David Ewers, Timon J Hartmann, Stefan Volkmann, Theresa Kungl, Robert Fledrich, Sandra Goebbels, Klaus-Armin Nave, Michael W Sereda

## Abstract

Duplication of the gene encoding the myelin protein PMP22 causes the hereditary neuropathy Charcot-Marie-Tooth disease 1A (CMT1A), characterized by hypomyelination of medium to large caliber peripheral axons. Conversely, haplo-insufficiency of *PMP22* leads to focal myelin overgrowth in hereditary neuropathy with liability to pressure palsies (HNPP). However, the molecular mechanisms of myelin growth regulation by PMP22 remain obscure. Here, we found that the major inhibitor of the myelin growth signaling pathway PI3K/Akt/mTOR, phosphatase and tensin homolog (PTEN) is increased in abundance in CMT1A and decreased in HNPP rodent models. Indeed, treatment of DRG co-cultures from HNPP mice with PI3K/Akt/mTOR pathway inhibitors reduced focal hypermyelination and, importantly, treatment of HNPP mice with the mTOR inhibitor Rapamycin improved motor behavior, increased compound muscle amplitudes (CMAP) and reduced tomacula formation in the peripheral nerve. In *Pmp22*^*tg*^ CMT1A mice, we uncovered that the differentiation defect of Schwann cells is independent from PI3K/Akt/mTOR activity, rendering the pathway insufficient as a therapy target on its own. Thus, while CMT1A pathogenesis is governed by dys-differentiation uncoupled from PI3K/Akt/mTOR signaling, targeting the pathway provides novel proof-of-principle for a therapeutic approach to HNPP.

## Introduction

Schwann cells wrap myelin around peripheral nerve axons for fast neural transmission^1^. Proper expression of the peripheral myelin protein of 22 kDa (PMP22), an integral constituent of the compact myelin sheath, is important for development and function of peripheral nerve fibers. The *PMP22* gene is located on chromosome 17p11.2 and mutations as well as alterations in the gene dosage are causative for a group of hereditary neuropathies affecting approximately 1 in 2500 humans^2,3^. Haplo-insufficiency of *PMP22* causes hereditary neuropathy with liability to pressure palsies (HNPP)^4^. Clinically, patients suffer from sensory loss as well as palsies and paresthesia upon mechanical stress on the nerve^5,6^. A hallmark of the HNPP disease is the formation of tomacula, extensive formation of myelin sheaths at cytoplasmic areas, such as paranodes and Schmidt-Lanterman incisures leading to deformed and constricted axons and subsequently demyelination^7-9^. Slowed nerve conduction velocity and conduction block can be observed at sites susceptible for compression while other regions are unaffected^10,11^. Complementarily, a duplication on chromosome 17 results in the overexpression of PMP22 and causes Charcot-Marie-Tooth disease type 1A (CMT1A)^12-15^. Patients suffer from a distally pronounced, slowly progressive muscle weakness and sensory symptoms that are accompanied by slowed nerve conduction velocity (NCV)^16^. Schwann cells from CMT1A patients and animal models show early dys-differentiation resulting in small hypermyelinated axons and later onion bulb formation^17,18^ and large caliber axons that are thinly or amyelinated along with reduced internodal length and secondary axonal loss^19^. Although the genetics of *PMP22* gene dosage diseases have been described more than 25 years ago, the molecular mechanisms causative for the abnormal myelination remain largely unknown and no therapy is available today.

Myelin growth is regulated by the PI3K/Akt/mTOR signaling pathway in the nervous system. We have shown previously that *Pmp22* dosage impacts PI3K/Akt/mTOR signaling^17^. Here, we asked whether PI3K/Akt/mTOR signaling provides a therapeutical target to treat the consequences of altered *Pmp22* gene-dosage. We found a correlation between PMP22 dosage and protein levels of the major inhibitor of the PI3K/Akt/mTOR pathway, phosphatase and tensin homolog (PTEN). In CMT1A rodent models, direct targeting of PTEN improved myelination only transiently without improving a differentiation defect which is absent in HNPP. However, we demonstrate that HNPP, as modelled in haplo-insufficient *Pmp22*^*+/-*^ mice, can be alleviated by pharmacologically targeting mTOR signaling.

## Results

### PMP22 dosage perturbs the abundance of the growth signaling inhibitor PTEN in animal models of CMT1A and HNPP

The major inhibitor of the PI3K/Akt/mTOR pathway is Phosphatase and Tensin homolog (PTEN), which counteracts PI3K by dephosphorylating PIP3 to PIP2, resulting in a decreased activity of Akt^20,21^. Ablation of *Pten* in Schwann cells thus leads to an overactivation of the PI3K/Akt/mTOR pathway and focal hypermyelination of axons^22,23^, a phenotype similar to HNPP. We tested mouse mutants of HNPP (*Pmp22*^*+/-*^) by Western Blot analysis and found, consistently through postnatal development, PTEN to be reduced in nerves lacking normal PMP22 levels (**Figure 1a**). In contrast, in *Pmp22* transgenic rats (*Pmp22*^*tg*^) that model CMT1A, the overexpression of PMP22 correlated with increased abundance of PTEN (**Figure 1c**). In all animals, PTEN was most abundant at the peak of myelination (P18), suggesting a role for PTEN in development (**Figure 1a, c**). Interestingly, we observed the same difference at the mRNA level suggesting the *Pmp22* levels affect *Pten* transcription (**Figure 1b, d**). We subsequently performed Western Blot analysis and found that PTEN is more abundant in peripheral nerve lysates than in myelin (**Figure 1e**). Indeed, immunohistochemical staining revealed PTEN expression in axons, Schwann cell nuclei and cytoplasmic non-compact myelin compartments including the bands of Cajal (**Figure 1f**).

**Figure 1:**
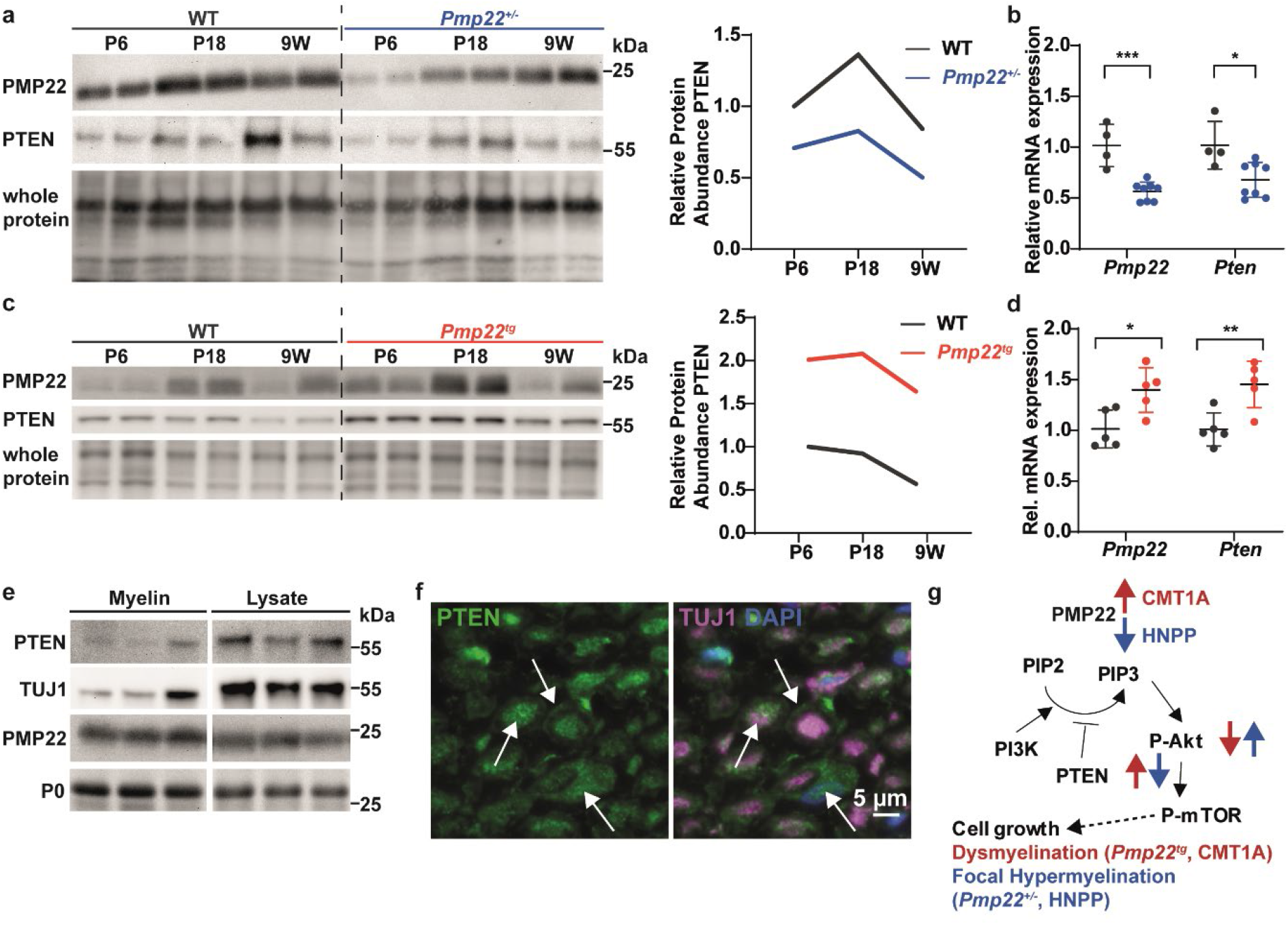
PTEN is *Pmp22* gene-dosage dependently altered in animal models of HNPP and CMT1A. **a** Western Blot analysis (left panel) showing decreased PTEN and PMP22 protein levels in sciatic nerve lysates of *Pmp22*^*+/-*^ mice at postnatal day 6 (P6), postnatal day 18 (P18) and 9 weeks of age compared to wildtype (WT) control (n = 2 per time point and group). Fast green whole protein staining was used as loading control for the quantification (right panel). **b** Quantitative RT-PCR analysis in tibial nerves shows decreased mRNA levels of *Pmp22* and *Pten* in *Pmp22*^*+/-*^ mice at P18. *Rplp0* and *Ppia* served as housekeeping genes. **c** Western Blot analysis (left panel) of PTEN and PMP22 protein levels in sciatic nerve lysates at P6, P18 and 9 weeks of age reveals increased protein levels in *Pmp22*^*tg*^ rats compared to WT control. Fast green whole protein staining served as loading control for the quantification (right panel). **d** Quantitative RT-PCR analysis in tibial nerves shows increased mRNA levels of *Pmp22* and *Pten* in *Pmp22*^*tg*^ rat tibial nerves at P18. *Rplp0* and *Ppia* served as housekeeping genes. **e** Immunoblot of WT P18 rat whole sciatic nerve lysate and purified myelin. PTEN and TUJ1 are enriched in the lysate while PMP22 and P0 are enriched in the myelin fraction. **f** Femoral nerve cross section of 9 weeks old WT rats shows PTEN (green) localization to the axon (magenta, TUJ1), nuclei (blue DAPI) and bands of Cajal (indicated by arrows). Scale bar is 5 µm. **g** Graphical overview of *Pmp22* gene-dosage dependent alterations in the PI3K/Akt/mTOR signaling pathway in animal models of CMT1A and HNPP. PMP22 overexpression leads to increased PTEN protein levels, reduced activation of the downstream PI3K/Akt/mTOR growth signaling pathway and subsequently demyelination (red). In contrast, PMP22 heterozygosity results in decreased PTEN levels, increased activation of the PI3K/Akt/mTOR signaling cascade and hypermyelination (blue). Statistical analysis was performed using student’s t-test, *p < 0.05,**p < 0.01, ***p < 0.001.

Taken together, this supports a working model in which *Pmp22* dosage inversely correlates with the activation of the PI3K/Akt/mTOR signaling via PTEN (**Figure 1g**). In *Pmp22*^*+/-*^ HNPP nerves the reduced levels of PMP22 enhance PI3K/Akt/mTOR signaling due to a partial loss of PTEN, leading to abnormal myelin growth while increased PMP22 levels reduce PI3K/Akt/mTOR activity in *Pmp22*^*tg*^ CMT1A nerves, leading to dysmyelination. The model prompted us to test the possibility that targeting the PI3K/Akt/mTOR pathway could be a therapeutic strategy.

### Targeting PI3K/Akt/mTOR signaling ameliorates myelin outfoldings in *Pmp22*^*+/-*^ HNPP co-cultures

To show *proof of principle*, we first studied Schwann cell dorsal root ganglia neuron (SC-DRG) co-cultures, derived from *Pmp22*^*+/-*^ mice, which were either treated with LY294002, a PI3K inhibitor, or with Rapamycin, an inhibitor of mTOR signaling (**Figure 2a**). *Pmp22*^*+/-*^ co-cultures showed myelin outfoldings, which were particularly prominent at paranodal regions (**Figure 2b**) and resembled tomacula, the histological hallmark of the HNPP disease. Immunocytochemical analysis revealed an approximately 35 % increase in the number of myelin segments with outfoldings in SC-DRG co-cultures from *Pmp22*^*+/-*^ mice, when tested two weeks after myelination was induced (**Figure 2c**). Subsequent treatment with both inhibitors, but not DMSO, for two weeks strongly decreased the number of segments harboring outfoldings to approximately 20 % (**Figure 2b, c**). Of note, the total number of myelinated segments was unaltered in mutant cultures (**Figure 2d**). Thus, inhibiting the activated PI3K/Akt/mTOR pathway in *Pmp22*^*+/-*^ Schwann cells rescued aberrant myelin growth *in vitro*.

**Figure 2:**
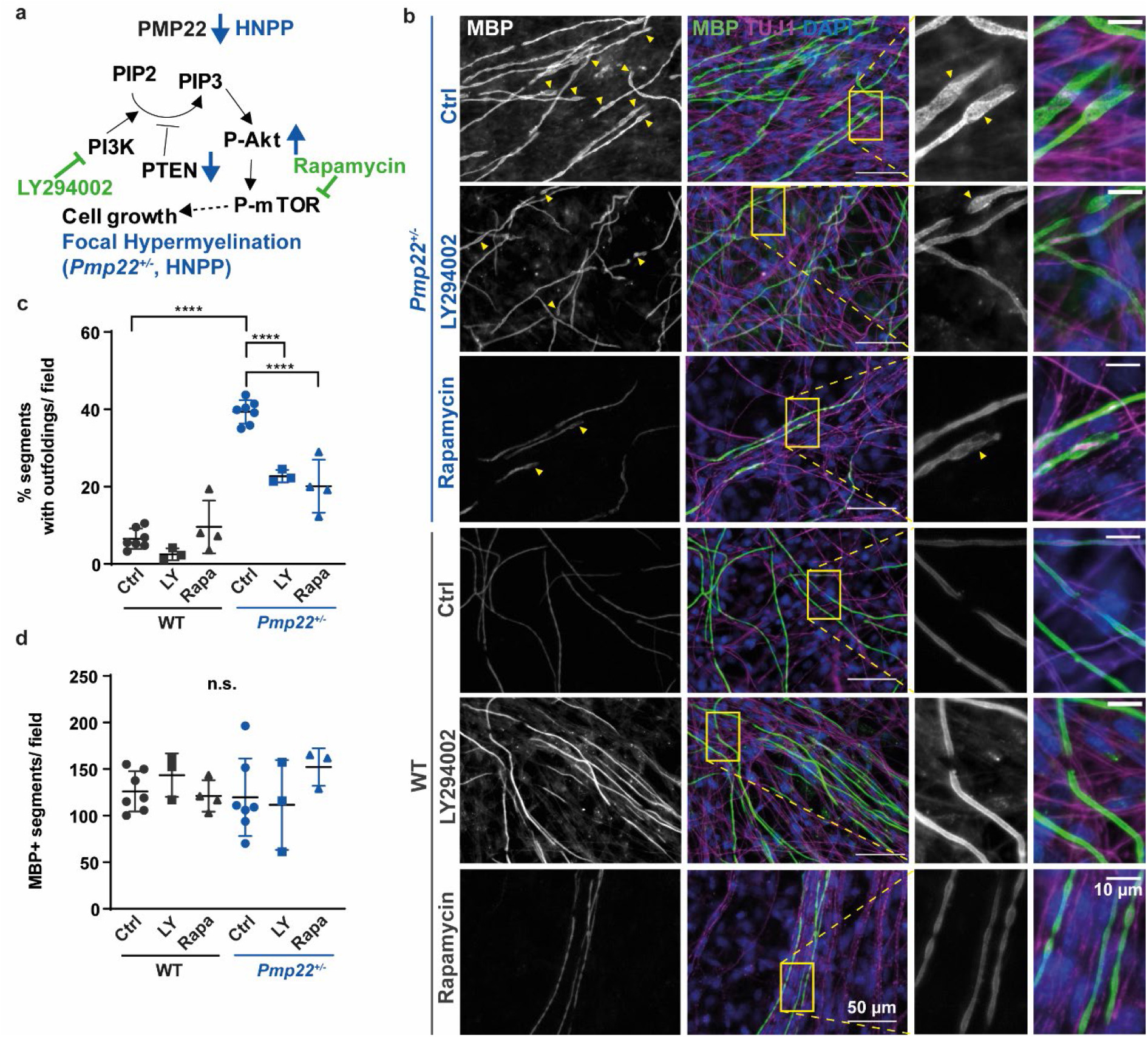
Inhibition of the PI3K/Akt/mTOR signaling pathway reduces myelin outfoldings in *Pmp22*^*+/-*^ co-cultures. **a** PI3K inhibitor LY294002 and mTOR inhibitor Rapamycin were used to counteract the upregulated PI3K/Akt/mTOR signaling pathway in *Pmp22*^*+/-*^ mice. **b** Example images displaying Schwann cell-dorsal root ganglia neuron co-cultures from wildtype (WT) and *Pmp22*^*+/-*^ (HNPP) mice 14 days after induction of myelination. Cells were treated with either DMSO as controls, 10 µM LY294002 or 20 nM Rapamycin. Fixed cells were stained for myelin basic protein (MBP, grey/green) to visualize myelin and beta tubulin III (TUJ1, magenta) for axons. Cell nuclei are stained by DAPI in blue. White arrowheads indicate myelin outfoldings. Scale bar is 50 μm (overview) and 10 μm (blow-up). **c** Quantification of **b** reveals an increased proportion of myelinated segments with outfoldings in *Pmp22*^*+/-*^ control co-cultures (circles) and a reduction after inhibition of PI3K (squares) or mTOR (triangles). n = 3-7 animals with n = 5 fields of view (500 × 500 µm) were quantified. **d** The mean number of myelinated segments is unaltered in control and treated *Pmp22*^*+/-*^ co-cultures. n = 3-7 animals with n = 5 fields of view (500 × 500 µm) were quantified. Mean numbers are displayed ± standard deviation. Groups were compared using one-way ANOVA with Sidak’s multiple comparison test (*p ≤ 0.05 **p ≤ 0.01, ***p ≤ 0.001, ****p ≤ 0.0001).

### Rapamycin treatment improves the phenotype of *Pmp22*^*+/-*^ HNPP mice

Following the promising *in vitro* findings, we performed a therapeutic *in vivo* trial by treating *Pmp22*^*+/-*^ mice with the mTOR inhibitor Rapamycin. Between postnatal days 21 and 148, mice were intraperitoneally injected with 5 mg Rapamycin per kilogram bodyweight twice per week (**Figure 3a**).

**Figure 3:**
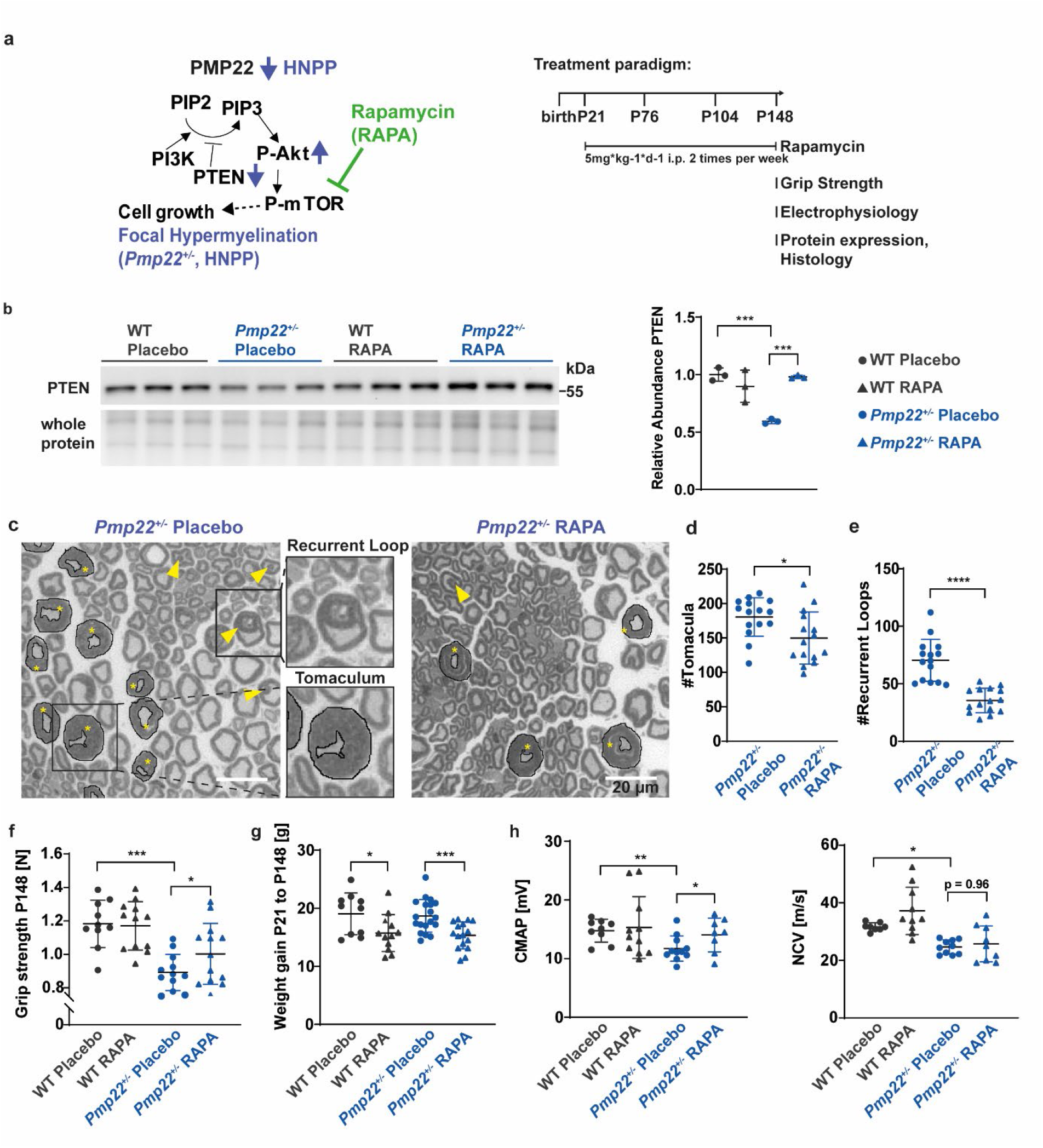
Rapamycin treatment in *Pmp22*^*+/-*^ mice ameliorates the disease phenotype. **a** *Pmp22*^*+/-*^ and wildtype (WT) control mice were injected i.p. with placebo solution or 5 mg Rapamycin per kg bodyweight two times per week from P21 until P148 to reduce mTOR activity. Grip strength analysis, electrophysiology, protein expression analysis and histology were performed at P148. **b** Western Blot analysis of PTEN protein (left panel) in whole sciatic nerve lysates of n = 3 animals per group. Quantification using whole protein staining as loading control shows increased PTEN protein levels after Rapamycin treatment in *Pmp22*^*+/-*^ mice (right panel). **c** Sciatic nerve semi-thin sections of *Pmp22*^*+/-*^ placebo and Rapamycin treated mice. Tomacula are encircled in black and marked with asterisks. Yellow arrowheads point to recurrent loops. Scale bar is 20 µm. **d** The number of Tomacula is decreased in whole sciatic nerves of Rapamycin treated *Pmp22*^*+/-*^mice (triangle) compared to placebo controls (circles) at P148, n = 15 animals per group. **e** The number of Recurrent Loops is decreased in whole sciatic nerves of Rapamycin treated *Pmp22*^*+/-*^ mice (triangle) compared to placebo controls (circles) at P148, n = 15 animals per group. **f** Forelimb grip strength is decreased in *Pmp22*^*+/-*^ placebo mice (n = 12, blue circles) compared to WT placebo mice (n = 10, grey circles), whereas Rapamycin treatment improves strength in *Pmp22*^*+/-*^ mice (n = 13, blue triangles) and does not affect WT mice (n = 12, grey triangles) at P148. **g** Rapamycin treated *Pmp22*^*+/-*^ and WT animals (triangles) gained less weight than Placebo controls (circles). Groups of n = 10 WT placebo, n = 12 WT Rapamycin, n = 19 *Pmp22*^*+/-*^ placebo and n = 16 *Pmp22*^*+/-*^ Rapamycin. **h** Electrophysiological analysis shows reduced compound muscle action potential amplitudes (CMAP, left panel) and nerve conduction velocities (NCV, right panel) in *Pmp22*^*+/-*^ placebo mice and increased CMAP after Rapamycin treatment. n = 8-9 WT placebo, n = 10-11 WT Rapamycin, n = 10-11 *Pmp22*^*+/-*^ placebo and n = 8-9 *Pmp22*^*+/-*^ Rapamycin. Mean numbers are displayed ± standard deviation. Statistical analysis for two groups was performed using student’s t-test. More than two groups were compared using one-way ANOVA with Sidak’s multiple comparison test (*p ≤ 0.05 **p ≤ 0.01, ***p ≤ 0.001, ****p ≤ 0.0001).

At the molecular level, the relative abundance of PTEN was restored in nerves of *Pmp22*^*+/-*^ mice that were treated with Rapamycin (**Figure 3b**). As this inhibitor acts on mTOR, downstream of PTEN, the upregulation of PTEN indicates a feedback signal of mTOR that needs to be explored. Histologically, nerves from *Pmp22*^*+/-*^ mice displayed the disease-typical formation of tomacula, i.e. areas of focal hypermyelination, recurrent loops, and myelin infoldings (**Figure 3c**). Quantification of these in sciatic nerves cross sections from Rapamycin treated animals revealed a significant decrease in the pathological hallmarks of HNPP (**Figure 3c-e**). Importantly, Rapamycin treatment significantly improved the grip strength of *Pmp22*^*+/-*^ mice, although they did not reach wildtype levels (**Figure 3f**). Whereas Rapamycin expectedly reduced the developmental weight gain of all animals, grip strength of wildtype mice was unaffected (**Figure 3f, g**). Electrophysiological measurements revealed lower compound muscle action potential (CMAP) amplitudes and slower nerve conduction velocities (NCV) in *Pmp22*^*+/-*^ mice. After Rapamycin treatment, amplitudes increased, whereas conduction velocities remained decreased (**Figure 3h**).

Taken together, reducing the abnormal activity of the PI3K/Akt/mTOR pathway in HNPP Schwann cells improves the phenotype in a mouse model of this disease, which makes the clinically available Rapamycin a promising drug candidate for treatment of human patients.

### Inhibiting PTEN improves myelination in SC-DRG co-culture model of CMT1A

*Pmp22* expression correlated with the abundance level of PTEN **(Figure 1)**. The data so far suggested that *Pmp22* overexpression in CMT1A causes neuropathy by *Pten*-dependent (partial) loss of PI3K/Akt/mTOR signaling in Schwann cells. To test this hypothesis, we generated *Pmp22*^*tg*^ CMT1A SC-DRG co-cultures and attempted to increase the signaling strength of the pathway by targeting PTEN. To this end we applied a small vanadium complex (VO-OHpic), a highly potent and specific inhibitor of PTEN that increases PtdIns(3,4,5)P3 and Akt phosphorylation^24^ (**Figure 4a**). Similar to the *in vivo* situation in patients and animal models^25^, SC-DRG co-cultures from a *Pmp22*^*tg*^ rodent model have fewer myelinated segments (**Figure 4b, c**). However, the number of myelinated segments was significantly increased in the presence of 500 nM VO-OHpic as compared to DMSO controls (**Figure 4b, c**). Surprisingly, a much stronger inhibition of PTEN (with 5 μM VO-OHpic) diminished myelination, independent of the *Pmp22* genotype (**Figure 4c, Supplementary Figure S1**). Similarly, the prolonged inhibition of PTEN with VO-OHpic (for 14 days) caused a dosage-dependent reduction in myelinated segments in wildtype co-cultures (**Figure 4c, Supplementary Figure S1**). As PTEN is expressed in both, neurons and glia, a negative effect of neuronal PTEN inhibition that interferes with myelination is likely.

**Figure 4:**
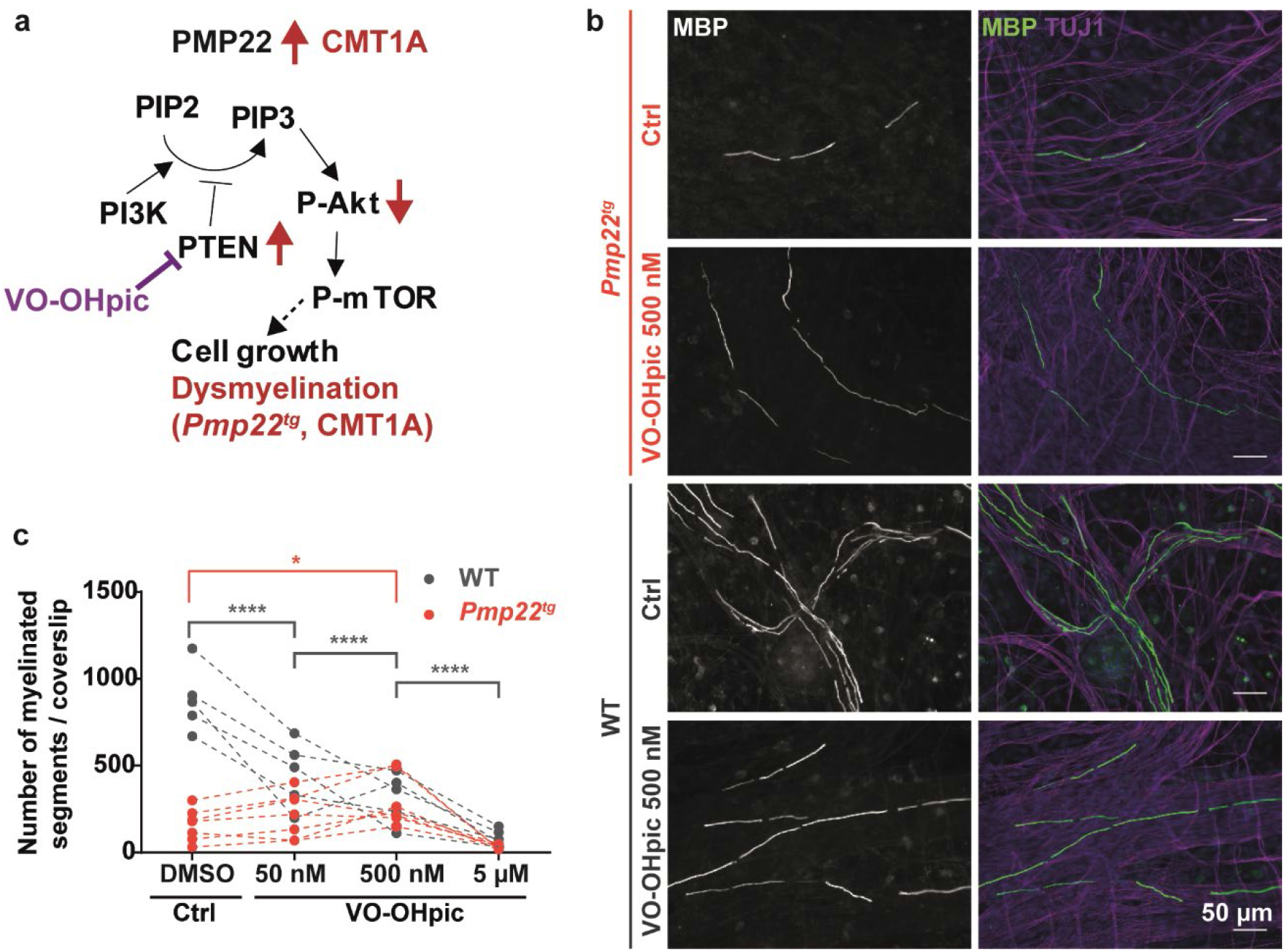
Inhibition of PTEN improves myelination in *Pmp22*^*tg*^ co-cultures *in vitro*. **a** PTEN inhibitor VO-OHpic was used to disinhibit the PI3K/Akt/mTOR signaling pathway in CMT1A. **b** Example images of Schwann cell (SC)-dorsal root ganglia neuron co-cultures from wildtype (WT) and *Pmp22*^*tg*^ rats treated with DMSO as controls (Ctrl), PTEN inhibitor VO-OHpic (500 nM). Cells were stained for myelin basic protein (MBP) as a marker for myelinated segments (grey/ green) and TUJ1 for neurons (magenta) as well as DAPI for cell nuclei (blue). **c** Quantification of **b** shows a dose-dependent decrease of myelinated segments in WT co-cultures treated with DMSO and different concentrations of VO-OHpic (50 nM, 500 nM, 5 µM) and an increase of myelinated segments in *Pmp22*^*tg*^ co-cultures with 500 nM VO-OHpic (WT n = 5, CMT1A n = 7 animals). Groups were compared using two-way ANOVA with Sidak’s multiple comparison test (*p ≤ 0.05 **p ≤ 0.01, ***p ≤ 0.001, ****p ≤ 0.0001).

### Targeting PTEN in Schwann cells of CMT1A mice

To study the impact of Schwann cell specific PTEN reduction in a CMT1A model *in vivo*, we applied a genetic approach (**Figure 5a**). We specifically reduced PTEN in Schwann cells (without affecting its neuronal expression) by crossbreeding *Pmp22*^*tg*^ mice with *Pten*^*flox*^ heterozygotes and *Dhh-cre* mice, yielding *PTEN*^*fl/+*^*Dhh*^*cre/+*^*PMP22*^*tg*^ experimental mutants (**Figure 5b**). Western blot analyses of sciatic nerve lysates confirmed a reduction of PTEN in the double mutants (**Figure 5c**) and revealed a corresponding activation of ribosomal protein S6, i.e. downstream of Akt and mTOR (**Figure 5d**). Strikingly, SC-DRG co-cultures derived from *PTEN*^*fl/+*^*Dhh*^*cre/+*^*Pmp22*^*tg*^ mice displayed an increased number of myelinated segments when compared to the *Pmp22*^*tg*^ single mutants (**Figure 5e, f**). In contrast, co-cultures from *PTEN*^*fl/+*^*Dhh*^*cre/+*^ mice showed no alteration in the number of myelinated segments (**Figure 5e, f**), suggesting that neuronal PTEN is also critical for myelination. Collectively, these data demonstrate that targeting PTEN in PMP22 overexpressing Schwann cells improves myelination, in line with our working model (**Figure 5a**).

**Figure 5:**
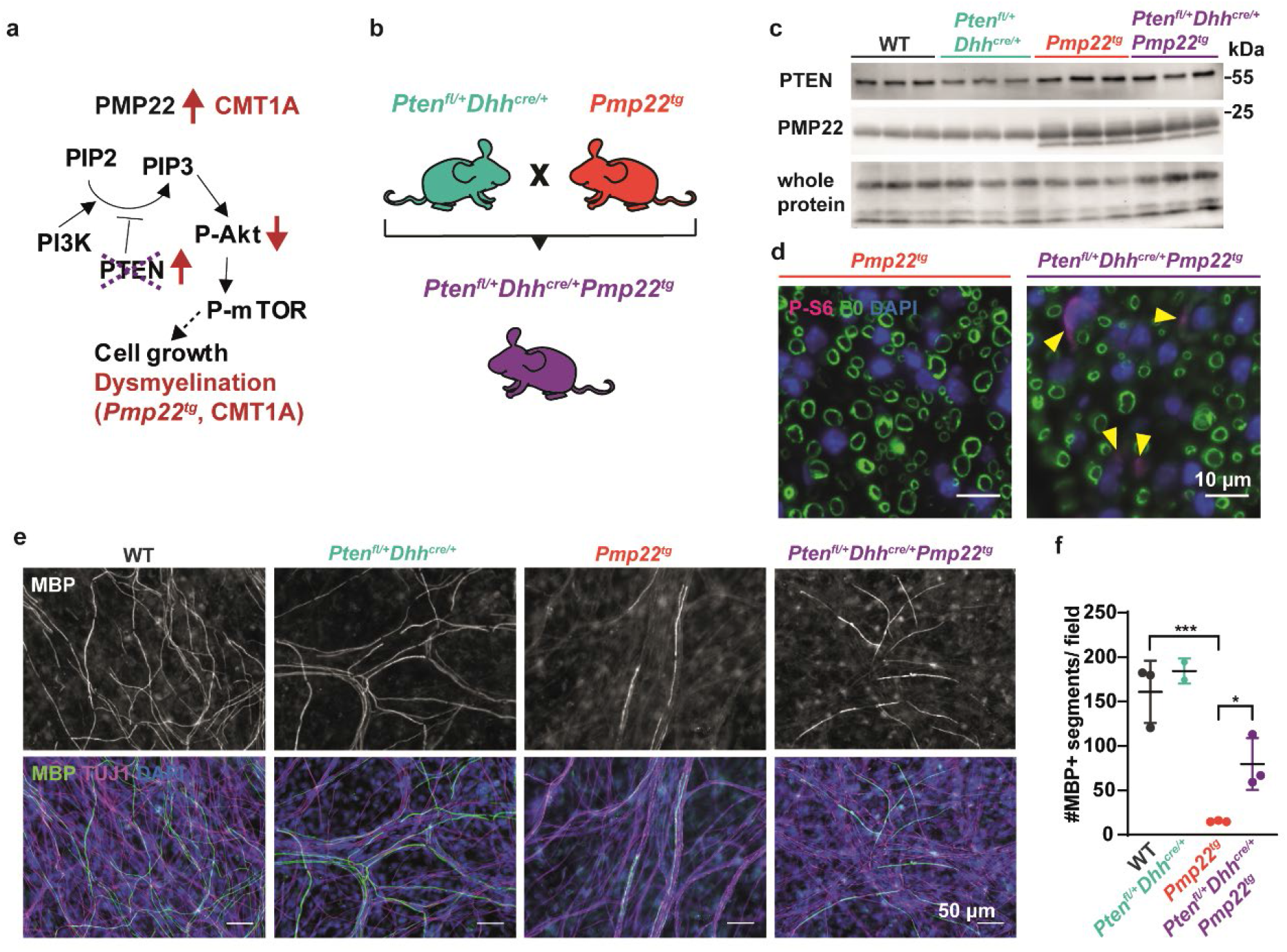
Genetic activation of the PI3K/Akt/mTOR signaling pathway increases myelination in *Pmp22*^*tg*^ co-cultures *in vitro*. **a** Crossing scheme of Schwann cell specific heterozygous Pten knockout mice (*PTEN*^*fl/fl*^*Dhh*^*cre/+*^) with CMT1A mice (*Pmp22*^*tg*^) to generate a heterozygous Pten knockout in CMT1A mice (*PTEN*^*fl/fl*^*Dhh*^*cre/+*^*Pmp22*^*tg*^). **b** In order to reduce Pten genetically in Schwann cells, heterozygous Pten floxed mice under the Dhh^cre^ driver (*Pten*^*fl/+*^*Dhh*^*cre/+*^) were crossbred with *Pmp22*^*tg*^ mice. **c** Western blot analysis shows PTEN and PMP22 protein amounts in whole sciatic nerve lysates from 16 weeks old WT, PTEN heterozygous knockout (*Pten*^*fl/+*^*Dhh*^*cre/+*^), CMT1A (*Pmp22*^*tg*^) and double mutant (*Pten*^*fl/+*^*Dhh*^*cre/+*^*Pmp22*^*tg*^) mice using whole protein staining as loading control. **d** Paraffin cross sections of femoral nerves from 18 days old *Pmp22*^*tg*^ and *Pten*^*fl/+*^*Dhh*^*cre/+*^*Pmp22*^*tg*^ mice show an increased signal for Phospho-S6 (magenta, indicated by yellow arrowheads). Myelin is visualized by P0 (green) and nuclei by DAPI (blue). Scale bar is 10 µm. **e** Representative example images of Schwann cell dorsal root ganglia neuron co-cultures 14 days after induction of myelination. Myelin basic protein (MBP) indicates myelinated segments (grey/green), TUJ1 neurons (magenta) and DAPI nuclei (blue). Scale bar is 50 µm. **f** Quantification of **e** shows increased numbers of myelinated segments in *Pten*^*fl/+*^*Dhh*^*cre/+*^*Pmp22*^*tg*^ compared to *Pten*^*fl/+*^*Dhh*^*cre/+*^ co-cultures. Shown are means of 5 fields of view (500 × 500 µm) for each animal (n = 2-3) ± standard deviation. Groups were compared using one-way ANOVA with Sidak’s multiple comparison test (*p ≤ 0.05, ***p ≤ 0.001).

We then investigated whether PTEN reduction alleviates the phenotype of *Pmp22*^*tg*^ mice also *in vivo. Pmp22*^*tg*^ mice had fewer myelinated axons and smaller axonal diameters when compared to their wildtype littermates at age P18 and 16 weeks (**Figure 6a, b, Supplement S2**)^26^. Plotting g-ratios against axon diameters revealed a shift towards hypermyelinated small caliber axons and hypomyelinated large caliber axons in *Pmp22*^*tg*^ mice (**Figure 6c, e**). In contrast, merely heterozygosity of *Pten* in Schwann cells did not significantly alter the number of myelinated axons, mean g-ratio or axon diameter (**Figure 6a, b Supplement S2**). Interestingly, *Pten* heterozygosity in *Pmp22*^*tg*^ mice increased the number of myelinated axons at postnatal day 18, but not in adult stages (**Figure 6b**), suggesting slightly advanced myelination. Similarly, we noted smaller g-ratios early in development (**Figure 6c, d**) but not at 16 weeks of age (**Figure 6e, f**). In adult mice, also the behavioral and electrophysiological readouts of *Pten* heterozygous *Pmp22*^*tg*^ mice were unaltered (**Figure 6g, h, Supplementary Figure S2**).

**Figure 6:**
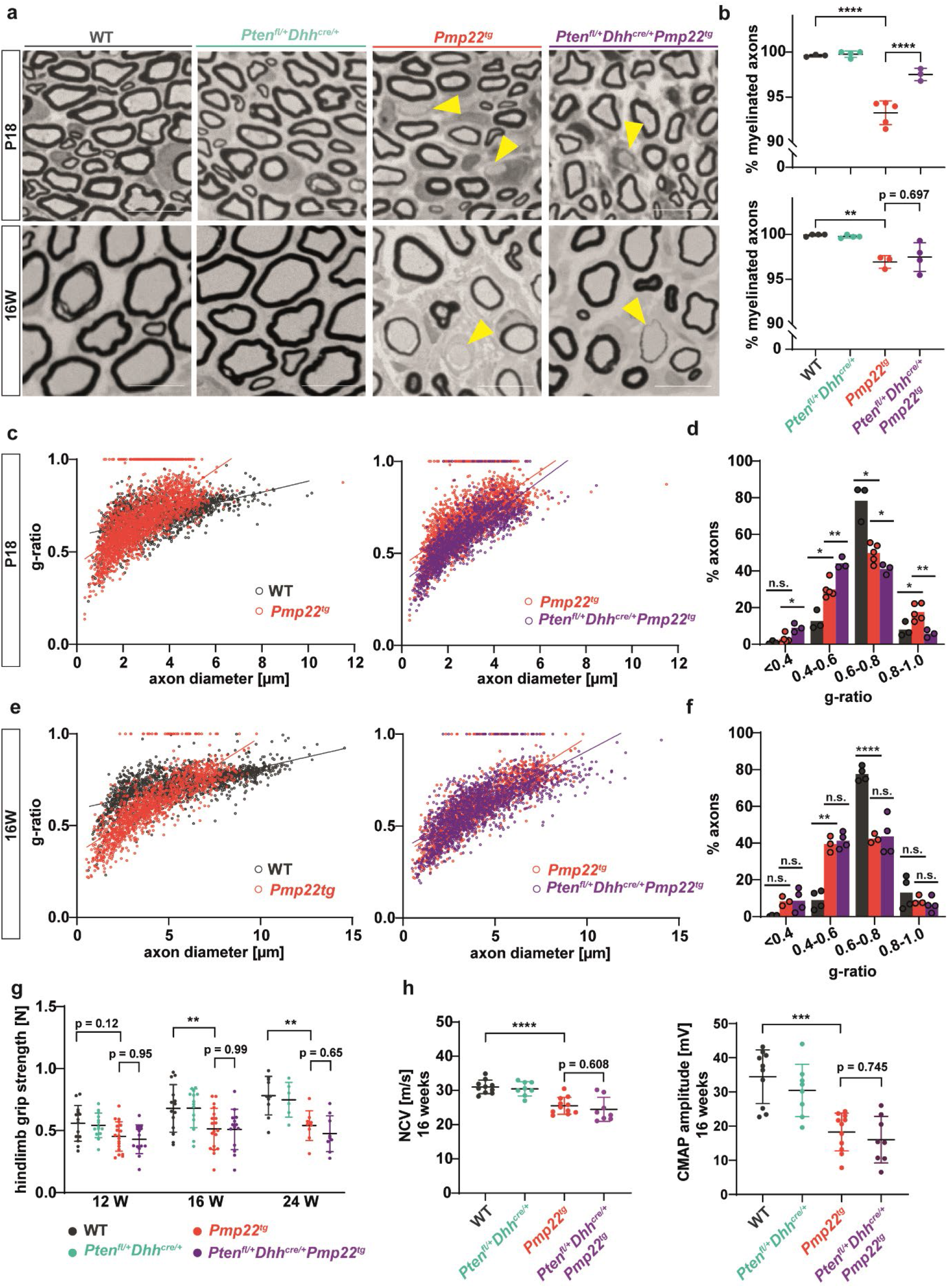
Reduction of PTEN in *Pmp22*^*tg*^ mice increases myelin growth early in development. **a** Example images of femoral nerve semi-thin sections from wildtype (WT), *Pten*^*fl/+*^*Dhh*^*cre/+*^, *Pmp22*^*tg*^ and *Pten*^*fl/+*^*Dhh*^*cre/+*^*Pmp22*^*tg*^ mice at P18 (upper panels) and 16 weeks of age (lower panels). Yellow arrowheads indicate amyelinated axons. Scale bar = 10 µm. **b** Quantification of **a** displays a decreased amount of myelinated axons in *Pmp22*^*tg*^ whole femoral nerves and an increase in *Pten*^*fl/+*^*Dhh*^*cre/+*^*Pmp22*^*tg*^ double mutants (upper panel) at P18 while numbers of myelinated axons are similarly decreased in *Pmp22*^*tg*^ and *Pten*^*fl/+*^*Dhh*^*cre/+*^*Pmp22*^*tg*^ mice at 16 weeks of age (lower panel). WT n = 3-4, *Pten*^*fl/+*^*Dhh*^*cre/+*^ n = 3-4, *Pmp22*^*tg*^ n = 3-5 and *Pten*^*fl/+*^*Dhh*^*cre/+*^*Pmp22*^*tg*^ n = 3-4 animals. **c** G-ratio plotted against axon diameter of WT (grey) and *Pmp22*^*tg*^ (red) mice (left panel) and *Pmp22*^*tg*^ and *Pten*^*fl/+*^*Dhh*^*cre/+*^*Pmp22*^*tg*^ (purple) mice (right panel) at P18. WT n = 3, *Pmp22*^*tg*^ n = 5 and *Pten*^*fl/+*^*Dhh*^*cre/+*^*Pmp22*^*tg*^ n = 3 animals. **d** *Pten*^*fl/+*^*Dhh*^*cre/+*^*Pmp22*^*tg*^ femoral nerves display more axons with small g-ratios (0.4-0.5) and less axons with bigger g-ratios (0.8-.0.9, 0.9-1.0) compared to *Pmp22*^*tg*^ nerves. WT n = 3, *Pmp22*^*tg*^ n = 5 and *Pten*^*fl/+*^*Dhh*^*cre/+*^*Pmp22*^*tg*^ n = 3 animals. Groups were compared using two-way ANOVA with Tukey’s multiple comparison test (*p ≤ 0.05, **p ≤ 0.01, ****p ≤ 0.0001). **e** G-ratio plotted against axon diameter of WT (grey) and *Pmp22*^*tg*^ (red) mice (left panel) and *Pmp22*^*tg*^ and *Pten*^*fl/+*^*Dhh*^*cre/+*^*Pmp22*^*tg*^ (purple) mice (right panel) at 16 weeks of age. WT n = 4, *Pmp22*^*tg*^ n = 3 and *Pten*^*fl/+*^*Dhh*^*cre/+*^*Pmp22*^*tg*^ n = 4 animals. **f** No alteration in the distribution of axons over the g-ratio is observed comparing femoral nerves from *Pmp22*^*tg*^ and *Pten*^*fl/+*^*Dhh*^*cre/+*^*Pmp22*^*tg*^ mice. WT n = 4, *Pmp22*^*tg*^ n = 3 and *Pten*^*fl/+*^*Dhh*^*cre/+*^*Pmp22*^*tg*^ n = 4 animals. Groups were compared using two-way ANOVA with Tukey’s multiple comparison test (*p ≤ 0.05, **p ≤ 0.01, ****p ≤ 0.0001). **g** The grip strength of hindlimbs is reduced in *Pmp22*^*tg*^ and *Pten*^*fl/+*^*Dhh*^*cre/+*^*Pmp22*^*tg*^ mice compared to WT controls and *Pten*^*fl/+*^*Dhh*^*cre/+*^ mice. Behavioral analysis was done at 12, 16 and 24 weeks of age. WT n = 10-14, *Pten*^*fl/+*^*Dhh*^*cre/+*^ n = 6-14, *Pmp22*^*tg*^ n = 9-19 and *Pten*^*fl/+*^*Dhh*^*cre/+*^*Pmp22*^*tg*^ n= 9-14 mice were analyzed. **h** Nerve conduction velocity (NCV, left panel) and Compound muscle action potential amplitudes (CMAP, right panel) are similarly slower in *Pmp22*^*tg*^ and *Pten*^*fl/+*^*Dhh*^*cre/+*^*Pmp22*^*tg*^ mice compared to WT controls. For electrophysiology measurements WT n = 10, *Pten*^*fl/+*^*Dhh*^*cre/+*^ n = 8, *Pmp22*^*tg*^ n = 11 and *Pten*^*fl/+*^*Dhh*^*cre/+*^*Pmp22*^*tg*^ n= 8 mice were analyzed. Means are displayed ± standard deviation. Statistical analysis was performed using one-way ANOVA with Sidak’s multiple comparison test (*p ≤ 0.05, **p ≤ 0.01, ***p ≤ 0.001, ****p ≤ 0.0001) if not indicated otherwise.

Thus, activating the PI3K/Akt/mTOR pathway in Schwann cells of *Pmp22*^*tg*^ mice *in vivo* by reducing its main inhibitor PTEN, partially prevented axons from being amyelinated and increased myelin sheath thickness but only transiently in early development with stronger hypermyelination of small caliber axons and less hypomyelination of large fibers (**Figure 6c**). In contrast, teased fiber preparations showed indistinguishably shortened internodes in both, *Pmp22*^*tg*^ and *PTEN*^*fl/+*^*Dhh*^*cre/+*^*Pmp22*^*tg*^ mice (**Supplementary Figure S2**). Thus, stimulating the PI3K/Akt/mTOR pathway in *Pmp22*^*tg*^ Schwann cells drives radial but not longitudinal myelin growth in early development.

### Schwann cell differentiation defects in CMT1A are not rescued by PTEN reduction

CMT1A Schwann cells display a differentiation defect^17^. To test whether PTEN has an impact on Schwann cell differentiation in *Pmp22*^*tg*^ mice, we compared the expression levels of ‘immaturity genes’ such as *Ngfr, Pou3f1* and *Sox2* but found no difference in *Pmp22* overexpressing mice when introducing *Pten* heterozygosity (**Figure 7a**). Importantly, these transcripts were never increased in nerves of *Pmp22*^*+/-*^ HNPP mice (**Figure 7b**). Similarly, the myelin sheath thickness of axons without tomacula in *Pmp22*^*+/-*^ mice was similar to wildtype controls at postnatal day 18, whereas *Pmp22*^*tg*^ CMT1A nerves displayed a characteristic shift of hypermyelinated small axons and thinly myelinated larger axons at that age (**Figure 7c)**. In our resulting working model, *Pmp22* heterozygosity causes HNPP, which is clearly mediated by a decrease in PTEN levels and the resulting increase in PI3K/Akt/mTOR signaling (**Figure 7d**). Tomacula formation was improved by inhibiting mTOR using Rapamycin. *Pmp22* overexpression in CMT1A causes the Schwann cell differentiation defect and an increase in PTEN levels with the corresponding reduction of PI3K/Akt/mTOR signaling in Schwann cells. In contrast to HNPP, reducing PTEN levels in CMT1A did not overcome the differentiation defect.

**Figure 7:**
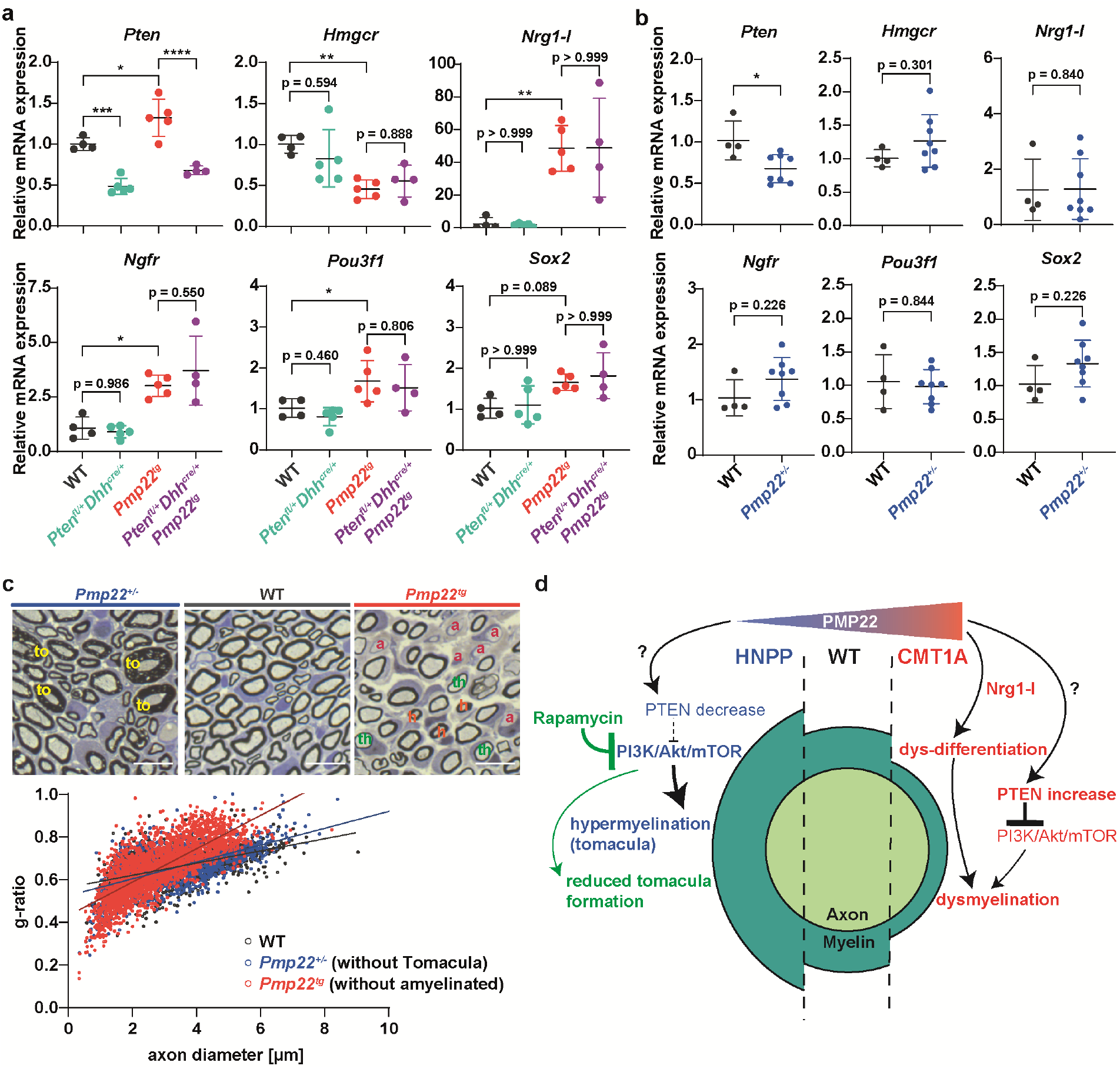
PMP22 overexpression specific differentiation defect is not rescued by PTEN reduction. **a** Quantitative RT-PCR from P18 tibial nerves of wildtype (WT) (n = 4), *Pten*^*fl/+*^*Dhh*^*cre/+*^ (n = 5), *Pmp22*^*tg*^ (n = 5) and *Pten*^*fl/+*^*Dhh*^*cre/+*^*Pmp22*^*tg*^ (n = 5) mice shows relative mRNA expression of *Pten, Hmgcr, Nrg1-I, Pou3f1, Ngfr*, and *Sox2. Rplp0* and *Ppia* were used as housekeeping genes. **b** Quantitative RT-PCR from P18 tibial nerves of WT (n = 4) and *Pmp22*^*+/-*^ (n = 8) mice shows relative mRNA expression of *Pten, Hmgcr, Nrg1-I, Pou3f1, Ngfr*, and *Sox2. Rplp0* and *Ppia* were used as housekeeping genes. Means are displayed ± standard deviation. **c** Semi-thin sections of P18 femoral nerves of *Pmp22*^*+/-*^ (left), WT (middle) and *Pmp22*^*tg*^ (right) mice. *Pmp22*^*+/-*^ nerves show myelin overgrowth, the so-called tomacula (to, yellow) and *Pmp22*^*tg*^ nerves are characterized by amyelinated (a, pink) and thinly myelinated (th, green) big axons as well as hypermyelinated (h, orange) small axons. Quantification shows g-ratios of *Pmp22*^*+/-*^ mice (blue, without tomacula) are similar to the WT (grey) distribution, while *Pmp22*^*tg*^ mice (red, without amyelinated) show the hypermyelination of small axons and demyelination of big axons. **d** In HNPP (*Pmp22* haplo-insufficiency), the PTEN levels are decreased leading to an increased activity of the PI3K/Akt/mTOR signaling pathway and focal hypermyelination (tomacula). The phenotype is ameliorated by inhibiting mTOR with Rapamycin and thus reducing tomacula formation. In CMT1A (*Pmp22* overexpression), Schwann cells show both a differentiation defect mediated by Nrg1-I and less activity of the PI3K/Akt/mTOR signaling pathway due to increased PTEN levels leading to a dysmyelination phenotype. Means are displayed ± standard deviation. Statistical analysis was performed using one-way ANOVA with Sidak’s multiple comparison test (*p ≤ 0.05, **p ≤ 0.01, ***p ≤ 0.001, ****p ≤ 0.0001).

## Discussion

*Pmp22* is a dosage sensitive gene which causes Charcot-Marie-Tooth disease type 1A when overexpressed and hereditary neuropathy with liability to pressure palsies in haplo-insufficiency. We have made the unexpected observation that *PMP22* gene dosage not only determines the expression level of this myelin protein in the peripheral nervous system, but also affects the abundance of PTEN in Schwann cells. The phosphatase PTEN degrades the signaling lipid phospho-inositide-3 phosphate (PIP3). Thus, in *Pmp22*^*tg*^ (CMT1A) nerve fibers the increased abundance of PTEN most likely explains the previously described decrease of PI3K/Akt/mTOR signaling^17^, underlying reduced myelin growth in CMT1A. Our proposed model of disease (**Figure 7d**) is further supported by the observations that ablating Akt or mTOR in Schwann cells leads to hypomyelination^27-29^, whereas the transgenic overexpression of Akt in these cells, similar to targeting PTEN directly, causes hypermyelination^22,23,30,31^. Indeed, depleting PTEN in Schwann cells causes an HNPP-like phenotype^22^ with focal myelin hypergrowth (tomacula). Strikingly, we could show that treatment of *Pmp22*^*+/-*^ (HNPP) mice with Rapamycin, a specific inhibitor mTOR and clinically approved drug, ameliorated the disease phenotype.

*PTEN* is a tumor suppressor gene that mediates growth arrest^32^, similar to PMP22’s initially described function as a growth arrest specific gene from fibroblasts^33^. In Schwann cells, PTEN interacts with discs large homolog 1 (Dlg1) at the adaxonal myelin membrane, where the downregulation of PI3K/Akt/mTOR signaling terminates myelination^27^. Interestingly, deletion of *Pten* in Schwann cells leads to a delayed onset of myelination early postnatally^23^. Similarly, in *Pmp22*^*+/-*^ mice the onset of myelination is delayed, whereas at later stages these mice exhibit the HNPP phenotype with tomacula formation^34^. Thus, PMP22 serves two functions that affect the timing of myelination. Early in development, PMP22 enables the timely onset of myelination, whereas in later development PMP22 puts a break on myelin growth. We note that also mTOR plays such a dual role, as increased mTOR activity in early development suppresses myelination (following radial sorting), whereas at later stages it is the loss of mTOR activity that interferes with continued myelination^23^. Thus, in CMT1A patients, the overexpression of PMP22 causes initially hypermyelination, most strikingly of small caliber axons, but a hypomyelination at later stages. Interestingly, upon strong PMP22 overexpression, Schwann cells cease to myelinate axons already right after sorting, i.e. before myelination has begun, as observed in *Pmp22* transgenic rats bred to homozygosity^35,36^.

The mechanisms that link the abundance of PMP22, a small cholesterol-binding tetraspan of the myelin sheath^37^, to that of PTEN are still unclear. By immunoelectron microscopy or co-immunoprecipitation, we failed to detect direct interactions between these proteins, i.e. in tissue extracts or when overexpressed in cultured cells (data not shown). This makes it highly likely that PMP22 stabilizes PTEN indirectly, for example by sequestration in membrane lipid rafts that are associated with PTEN interacting lipids, such as PIP2^38^. Theoretically, PTEN could also increase in abundance by binding to scaffolding proteins in PMP22-dependent membrane microdomains^39^.

The therapeutic effect of the mTOR inhibitor Rapamycin in a mouse model of HNPP suggests that Schwann cell mTOR signaling must be considered a pharmacological target in human HNPP for which no therapy is available^40^. Although generally considered a mild neuropathy, the clinical spectrum covers more severe disease courses, including recurrent positional short-term sensory symptoms, progressive mononeuropathy, Charcot–Marie–Tooth disease-like polyneuropathy, chronic sensory polyneuropathy, and chronic inflammatory demyelinating polyneuropathy-like, recurrent subacute polyneuropathy^5^. The range of duration of symptoms can reach from 1 week to 50 years^41^. In the cases of non-recovery, a local transdermal application of Rapamycin could provide a therapeutic strategy for affected peripheral nerves in the absence of systemic side effects. Rapamycin is a clinically approved drug (commercial name ‘Sirolimus’) that is mostly used as an immunosuppressive after organ transplantation and currently investigated in cancer therapy^42^. We note that systemic application of Sirolimus was generally well tolerated in a phase 2b study in the majority of patients with inclusion body myositis, a rare non-lethal neuromuscular disease^43^. Furthermore, activation of mTOR is observed in rodent models of other peripheral tomaculous neuropathies, e.g. following mutations in the phosphoinositide lipid phosphatases myotubularin-related protein 2 (MTMR2) and 13 (MTMR13) genes ^44-47^, thereby providing a common target for therapy^48^.

Why is there this striking therapeutic effect of correcting PI3K/Akt/mTOR signaling in *Pmp22*^*+/-*^ mice but no such effect in *Pmp22*^*tg*^ mice that model CMT1A? One possible explanation is that in CMT1A a mere 50% reduction of PTEN (*PTEN*^*fl/+*^*Pmp22*^*tg*^*Dhh*^*cre/+*^) stimulates mTOR but is not sufficient. In contrast, even the complete loss of PTEN from PMP22 overexpressing CMT1A Schwann cells (*PTEN*^*fl/fl*^*Pmp22*^*tg*^*Dhh*^*cre/+*^) leads at best to the formation of aberrant myelin profiles (**Supplementary Figure S3**). We observed that activating Akt by PTEN reduction increased myelin sheath thickness transiently at postnatal day 18 in CMT1A mice. However, not only large caliber axons were effected but also the already hypermyelinated smaller sized axons. Previously we could show that the persistent glial induction of Neuregulin 1 type I accounts for the hypermyelination of small caliber axons in CMT1A^18^. Increased levels of glial Neuregulin 1 type I are observed after peripheral nerve injury and in models of dysmyelinating neuropathies such as CMT1A and CMT1B^18,49^. Here we found increased expression of Neuregulin 1 type I and dysdifferentiation markers in nerves of *Pmp22*^*tg*^ (CMT1A) mice but not *Pmp22*^*+/-*^ (HNPP) mice. Moreover, PTEN reduction in *Pmp22*^*tg*^ mice did not alter the expression of those markers. Altogether we conclude that the Neuregulin 1 type I mediated differentiation defect is specific for the PMP22 overexpressing situation and independent from the PI3K/Akt/mTOR signaling pathway. This may explain the positive effects of counteracting the PI3K/Akt/mTOR pathway in *Pmp22*^*+/-*^ mice, whereas PTEN reduction slightly increased myelin growth early in development but did not overcome the differentiation defect as the dominating disease factor in *Pmp22*^*tg*^ mice.

In conclusion, our findings identify that PMP22 has a central role in timing myelination via the PTEN-PI3K/Akt/mTOR signaling axis. However, the dysdifferentiation phenotype observed upon PMP22 overexpression in CMT1A is uncoupled from the PI3K/Akt/mTOR signaling pathway. Future research aiming at effective therapy strategies may focus on a combination of targeting the differentiation defect and the dysregulated PI3K/Akt/mTOR pathway. Nonetheless such a differentiation defect is not observed upon haplo-insufficiency of *Pmp22* in HNPP, and counteracting the dysregulated signaling pathway by local administration of mTOR inhibitors provides a novel therapy strategy for patients.

## Material and Methods

### Transgenic rats and mice

The generation and genotyping of *Pmp22* transgenic mice (*Tg(PMP22)C61Clh*)^26^, *Pmp22* transgenic rats (*SD-Tg(Pmp22)Kan*)^35^, *Pmp22*^*+/-*^ mice (*Pmp22tm1Ueli*)^34^, *Pten-flox* mice (*Ptentm1HWu/J*)^50^ and the Dhh-Cre driver line (*FVB(Cg)-Tg(Dhh-cre)1Mejr/J*)^51^ have previously been described. In short, for genotyping, genomic DNA was isolated from ear or tail biopsies by incubation in Gitschier buffer with TritonX-100 and proteinase K for a minimum of 2 h at 55 °C following heat inactivation of proteinase K at 90 °C for 10 minutes. Genotyping primers used for routine genotyping are listed in supplementary table 1.

All animal experiments were conducted according to the Lower Saxony State regulations for animal experimentation in Germany as approved by the Niedersächsische Landesamt für Verbraucherschutz und Lebensmittelsicherheit (LAVES) and in compliance with the guidelines of the Max Planck Institute of Experimental Medicine.

Inclusion and exclusion criteria were pre-established. Animals were randomly included into the experiment according to genotyping results, age and weight and independent of their sex. Animals were excluded prior to experiments in case of impaired health conditions or weight differences to the average group of more than 10 %. During or after the experiment exclusion criteria comprise impaired health condition of single animals not attributed to genotype or experiment (according to veterinary) or weight loss > 10 % of the average group. No animals had to be excluded due to illness/ weight loss > 10 % of the average group. Exclusion criteria regarding the outcome assessment were determined with an appropriate statistical test, the Grubbs’ test (or ESD method) using the statistic software Graph Pad (Prism). Animal experiments (phenotype analysis, electrophysiology and histology) were conducted in a single blinded fashion towards the investigator. Selection of animal samples out of different experimental groups for molecular biology/ histology/ biochemistry was performed randomly and in a blinded fashion.

### Rapamycin therapy

Rapamycin (LC laboratories) was dissolved in vehicle solution containing 5 % polyethyleneglycol 400, 5 % Tween 80 and 4 % ethyl alcohol. HNPP and control mice were i.p. injected with 5 mg Rapamycin per kg bodyweight two times per week from postnatal day 21 until postnatal day 148 with either Rapamycin or placebo (vehicle solution). The weight was continuously controlled and animals were subjected to motor phenotyping tests and electrophysiology at the end of the study.

### Motor phenotyping

The same investigator per experimental group who was blinded towards the genotype performed all phenotyping analyses. Motor performance was assessed by standardized grip strength test and elevated beam test. To assess grip strength, the animals were hold by their tail and placed with their forelimbs on a horizontal bar. By gently pulling the animal away, the maximum force was measured in a connected gauge (FMI-210B2 Force Gauge, Alluris). To assess hind limb grip strength, the animal’s forelimbs were supported and their hindlimbs placed on the bar. Again, the animal was retracted from the bar and the gauge detected the maximum force applied. All measurements were repeated seven times per animal and the mean calculated. In between fore- and hindlimb grip strength analysis, the animals had a minimum break of 10 minutes. The elevated beam is an 80 cm long, 14 mm wide bar approximately 60 cm above the ground. The animals were habituated to the beam one day prior to the experimental day. The animals walked the elevated beam for three times and the mean walking time and number of slips was calculated.

### Electrophysiology

For anesthesia, mice were injected i.p. with 6 mg Ketamine (Bayer Vital) and 90mg Xylazine (WDT) per kg bodyweight. When no toe reflexes were observed anymore, electrophysiological measurements on the sciatic nerve and tail were performed (Evidence 3102evo, Schreiber und Tholen). Therefore, needle electrodes were subcutaneously applied close to the sciatic notch (proximal stimulation) and in close proximity to the ankle (distal stimulation). Motor recording electrodes were inserted in the small muscle on the plantar surface. After proximal and distal supramaximal stimulation the compound muscle action potential (CMAP) was recorded. The distance between the stimulation sites [m] divided by the difference of latencies [s] allowed to calculate the motor nerve conduction velocity (mNCV). For sensory measurements in the tail, needle electrodes were applied close to the tip of the tail as stimulation electrodes and electrodes in the proximal tail branch served as recording electrodes. Averaged compound sensory nerve potentials were measured after stimulation with 5 mA. According to the distance of electrodes the sensory nerve conduction velocity (sNCV) was calculated.

### Histology

Peripheral nerves were fixed in 2.5 % glutaraldehyde and 4 % PFA in 0.1 M phosphate buffer for 1 week. Then, they were contrasted with osmium tetraoxide and embedded in epoxy resin (Serva) and 0.5 µm semi-thin sections were cut (Leica RM 2155) using a diamond knife (Histo HI 4317, Diatome). Afterwards sections were stained according to Gallyas^52^ and with Methylene blue/ Azur II for 1 min. Analysis was performed on total nerve cross sections using FIJI (NIH).

### Cell culture

Schwann cell (SC)-dorsal root ganglia (DRG) co-cultures were prepared from either E15.5 rat or E13.5 mouse embryos according to standard procedure^53^. After DRGs were digested with 0.25 % Trypsin (Invitrogen) at 37 °C for 45 minutes, the reaction was stopped by adding deactivated fetal calf serum (FCS; HyClone) and basic medium (1 % Penicillin/ Streptomycin (Lonza), 10 % FCS, 50 ng/ml nerve growth factor (NGF; Alomone Labs) in minimum essential medium (MEM; Gibco)) cells were plated on collagen-coated coverslips. On day 7, myelination was induced by culturing the cells in basic medium with 50 ng/ml ascorbic acid (AA; Sigma). For PTEN inhibition in *Pmp22*^*tg*^ cultures, cells were treated with 1 % DMSO (Sigma) as a control and with the PTEN inhibitor VO-OHpic^24^ (Sigma) at 50 nM, 500 nM and 5 μM, respectively. In *Pmp22*^*+/-*^ cultures, cells were treated with either 1 % DMSO as control, 20 nM mTOR inhibitor Rapamycin (LC Laboratories) or 10 µM PI3K inhibitor LY294002 (Cell Signaling, #9901). Medium was changed every 2-3 days for two weeks.

### Immunocytochemistry

Cells from SC-DRG co-cultures were fixed in 4 % paraformaldehyde (PFA) in 1x PBS for 10 minutes and then permeabilized in a mixture of 95 % ice-cold methanol and 5 % acetone at -20 °C for 5 minutes. Afterwards cells were incubated in blocking solution (2 % horse serum, 2 % bovine serum albumin (BSA), 0.1 % porcine gelantine) for 1 hour at room temperature before incubation in primary antibodies (mouse anti-MBP 1:500 (Covance), rabbit anti-TUJ1 1:250 (Covance)) over night at 4 °C. The next day, cells were washed in 1x PBS for three times and incubated in secondary antibodies (Alexa 488 donkey anti mouse 1:1000 (Invitrogen) and Alexa 568 donkey anti rabbit 1:1000 (Invitrogen) diluted in blocking solution with 0.2 µg/µl 4’,6’-diamidino-2-phenylindole (DAPI; Sigma) for 1 hour at room temperature. Following a three washing steps in PBS cells were mounted on slides in Mowiol mounting solution (9.6 % Mowiol (Sigma), 24 % Glycerol, 0.1 M Tris-HCl). Fluorescent images were taken using a Axiophot Observer Z (Zeiss) with a Colibri light source (Zeiss) and MRM camera (Zeiss). Acquisition and processing of the images was carried out using Zen2.6 blue software (Zeiss), FIJI (NIH) and Illustrator 2020 (Adobe).

### Immunohistochemistry

For immunohistochemistry analysis of cross sections, femoral nerves were dissected and immersion fixed in 4 % PFA for 24 hours and imbedded in paraffin. 5 µm cross sections were deparaffinized using a standard xylol and ethanol series. After target retrieval was induced by heating in citrate buffer samples were blocked in 10 % BSA and 20 % goat serum in PBS for 20 minutes. Incubation with primary antibodies (rabbit anti PTEN 1:50 (CST, #9188), mouse anti TUJ1 1:250 (Covance), mouse anti P0 1:200 (provided by J.J. Archelos^54^), rabbit anti Phospho-S6 (CST, #4858)) in PBS was carried out over night at 4 °C. To quantify the internodal length, teased fibers were prepared from P18 or 16 weeks old sciatic nerves. Following dissection, connective tissue was removed and nerves teased into single fibers using fine forceps, after 15 minutes of drying fibers were fixed in 4 % PFA for 5 minutes and permeabilized in ice-cold 95 % methanol and 5 % acetone for another five minutes. Furthermore, fibers were three times washed in PBS and incubated in blocking solution (10 % horse serum, 1 % BSA, 0.025 % Triton-X-100 in PBS) for 1 hour at room temperature. After incubation with primary antibodies (mouse anti MAG 1:50 (Chemicon clone 513), rabbit anti NaV1.6 1:250 (Alomone Labs #ASC-009)) in blocking solution, three washing steps in PBS followed. Secondary antibody (donkey anti mouse Alexa 488 1:1000, donkey anti rabbit Alexa 568 1:1000 (Invitrogen)) incubation was proceeded in the same way in cross sections and teased fibers, they were incubated with 0.2 µg/µl 4’,6’-diamidino-2-phenylindole (DAPI; Sigma) for 1 hour at room temperature. After three washing steps in PBS cells were mounted on slides in Mowiol mounting solution (9.6 % Mowiol (Sigma), 24 % Glycerol, 0.1 M Tris-HCl) and fluorescent images were obtained using a Axiophot Observer Z (Zeiss) with a Colibri light source (Zeiss) and MRM camera (Zeiss). Acquisition, processing and analysis of the images was carried out using Zen2.6 blue software (Zeiss), FIJI (NIH) and Illustrator 2020 (Adobe).

### Myelin purification

Myelin purification was performed as previously described^55^. In short, 6 dissected sciatic nerves of rats at postnatal day 18 were homogenized in 0.27 M sucrose (PreCellys, 6000 rpm, 2 × 15 s). Part of the homogenate was kept as ‘lysate control’ and the remaining homogenate was layered over 0.83 M sucrose in centrifugation tubes and centrifuged at 75000 x *g* for 30 min at 4 °C. The interface between the sucrose gradients (Myelin enriched fraction) was carefully transferred to a new centrifugation tube with 20 mM Tris-Cl buffer and again centrifuged at 75000 x *g* for 15 min at 4 °C. The pellet was resuspended in 20 mM Tris-Cl buffer solution and centrifuged at 12000 x *g* for 15 min. The final pellet (purified myelin) was taken up in 20 mM Tris-Cl buffer and further processed for protein analysis.

### Protein analysis

Sciatic nerves were homogenized using a Precellys24 homogenizer (VWR) in sucrose lysis buffer (270 nM sucrose, 10 mM Tris-HCl, 1 mM NaHCO_3_, 1 mM MgCl_2_, protease inhibitor (cOmplete Mini, Roche), phosphatase inhibitor (PhosphoStop, Roche)) and dissolved in loading buffer (40 % glycine [w/v], 240 mM Tris-HCl, 8 % SDS [w/v], 0.04 % bromophenol blue [w/v], 1 mM DTT). 15 µg protein was separated by electrophoresis using 4-16 % polyacrylamide gels (Mini-PROTEAN^®^TGX™, Biorad) and PageRuler Plus Prestained Protein Ladder (Thermo Fisher) as loading and size control. Afterwards, proteins were transferred to a methanol-activated Amersham Hybond PVDF membrane (GE Healthcare) at 100 V for 1 h at room temperature using Mini Trans-Blot Cell system (Biorad). Fast Green staining was used to detect the whole protein content on the membrane. Shortly, membrane were rinsed in water to remove transfer buffer residues, incubated in Fast Green staining solution (0.5 % Fast Green [w/v] (Sigma), 30 % methanol, 6.7 % glacial acetic acid) for 5 minutes, following a two times washing step in washing solution (30 % methanol, 6.7 % glacial acetic acid) before fluorescent imaging on an Intas ECL Chemostar (INTAS Science Imaging Instruments). After blocking in BSA-TBST (5 % BSA [w/v], 25 mM Tris, 75 mM NaCl, 5 % Tween 20 [w/v]) for 1 hour at room temperature, Western Blots were incubated overnight at 4 °C in primary antibodies against PMP22 (1:1000, Assay Biotech), PTEN (1:1000, CST #9188), TUJ1 (1:1000, Covance), P0 (1:000, J.J. Archelos^54^). After 3-5 washing steps in TBST, membranes were incubated in HRP secondary antibodies (1:5000, Dianova, 45 min room temperature) against the respective species and developed using Western Lightning Plus-ECL-Kit (Perkin Elmer) and the Intas Chemostar.

### RNA analysis

Total RNA was extracted from tibial nerves (without epineurium) using QIAzol lysis reagent and was purified according to manufacturer’s instructions using RNeasy Kit (Qiagen). The concentration and purity of the RNA was determined with the ratio of absorption at 260/280 nm using NanoDrop 2000 Spectraphotometer. Following, cDNA was synthesized from total RNA using poly-Thymin and random nonamer primers and Superscript III RNase H reverse transcriptase (Invitrogen). Quantitative real time PCR was carried out using the GoTaq qPCR Master Mix (Promega) and LC480 detection system (Roche). Reactions were carried out in four replicates. The cycle threshold (Ct) value was calculated using LC480 software (Roche) and mean values were normalized as fold changes to the control group. As internal standards peptidyl isomerase A (*Ppia*) and ribosomal protein large P0 (*Rplp0*) were used. Primer sequences are listed in supplementary table 2.

### Statistical Analysis

All data are presented as mean ± standard deviation unless indicated otherwise. Data processing and statistical analysis was performed using MS Excel 2016 and GraphPad Prism 8. Statistical tests are indicated in the figure legends. In short, two groups were compared using Student’s t-test. More than two groups were compared using one-way ANOVA with an appropriate post test. For comparing more than two groups for more than one time point (longitudinal analysis) the two-way ANOVA with appropriate post test was used. Statistical differences are marked by stars when *p < 0.05, **p < 0.01, ***p < 0.001 and ****p < 0.0001).

## Acknowledgements

We thank C. Huxley (Imperial College School of Medicine, London) for providing *Pmp22*^*tg*^ mice, D. Meijer (University Edinburgh) for *Dhh*^*Cre*^ mice and U. Suter (ETH Zürich) for *Pmp22*^*+/-*^ mice. Further we want to acknowledge U. Bode, B. Veith, C. Wieczorek, T. Hoffmeister, S. Schulze, T. Ruhwedel and W. Möbius (MPI of Experimental Medicine, Göttingen) for their excellent technical support. M. W. Sereda is supported by the German Ministry of Education and Research (BMBF, CMT-NET, FKZ: 01GM1511C and CMT-NRG, FKZ: 01GM1605), the German Research Foundation (DFG, SE 1944/3-1 with K. A Nave) and the Volkswagen Foundation (with D. Ewers). M. W. Sereda heads the Neurogenetics clinic and co-heads the NM-centre at UMG Göttingen. He is member of the Inherited Neuropathy Consortium RDCRN and of the European Reference Network for Rare Neuromuscular Diseases (ERN EURO-NMD).

## Author contributions

D.K., D.E., S.G., K.A.N. and M.W.S. designed the study. D.K. planned and performed experiments, analyzed the data and wrote the manuscript. D.E. performed myelin purification and contributed to the manuscript. T.J.H. and S.V. contributed to behavior experiments and histological analysis. T.K. and R.F. performed electrophysiological analysis and contributed to the manuscript. S.G., K.A.N. and M.W.S. supervised the study, contributed to discussion and the manuscript.

## Competing financial interest

The authors declare no competing interests.

## Supplementary data

**Supplementary figure S1:**
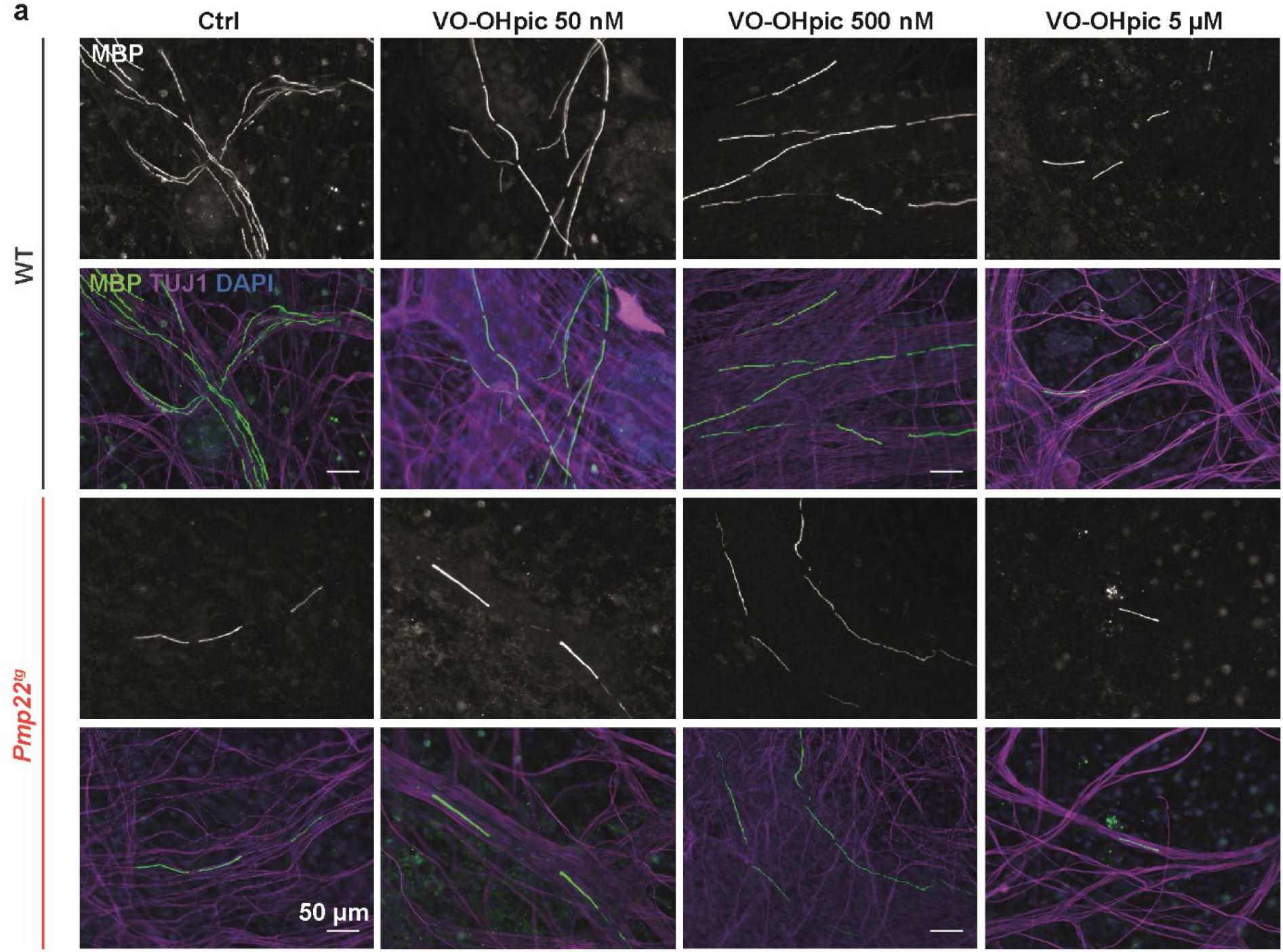
Dose-dependent response of myelination upon PTEN inhibition in *Pmp22*^*tg*^ co-cultures *in vitro*. **a** Example images of SC-DRG co-cultures from wildtype (WT) and *Pmp22*^*tg*^ rats, treated with different dosages of the PTEN inhibitor VO-OHpic. The number of myelinated segments (MBP; grey/green) decreases in WT cultures with increasing inhibitor dosage. In *Pmp22*^*tg*^ co-cultures an increase is observed up to 500 nM VO-OHpic but a decrease with 5 µM VO-OHpic. Scale bar is 50 µm.

**Supplementary figure S2:**
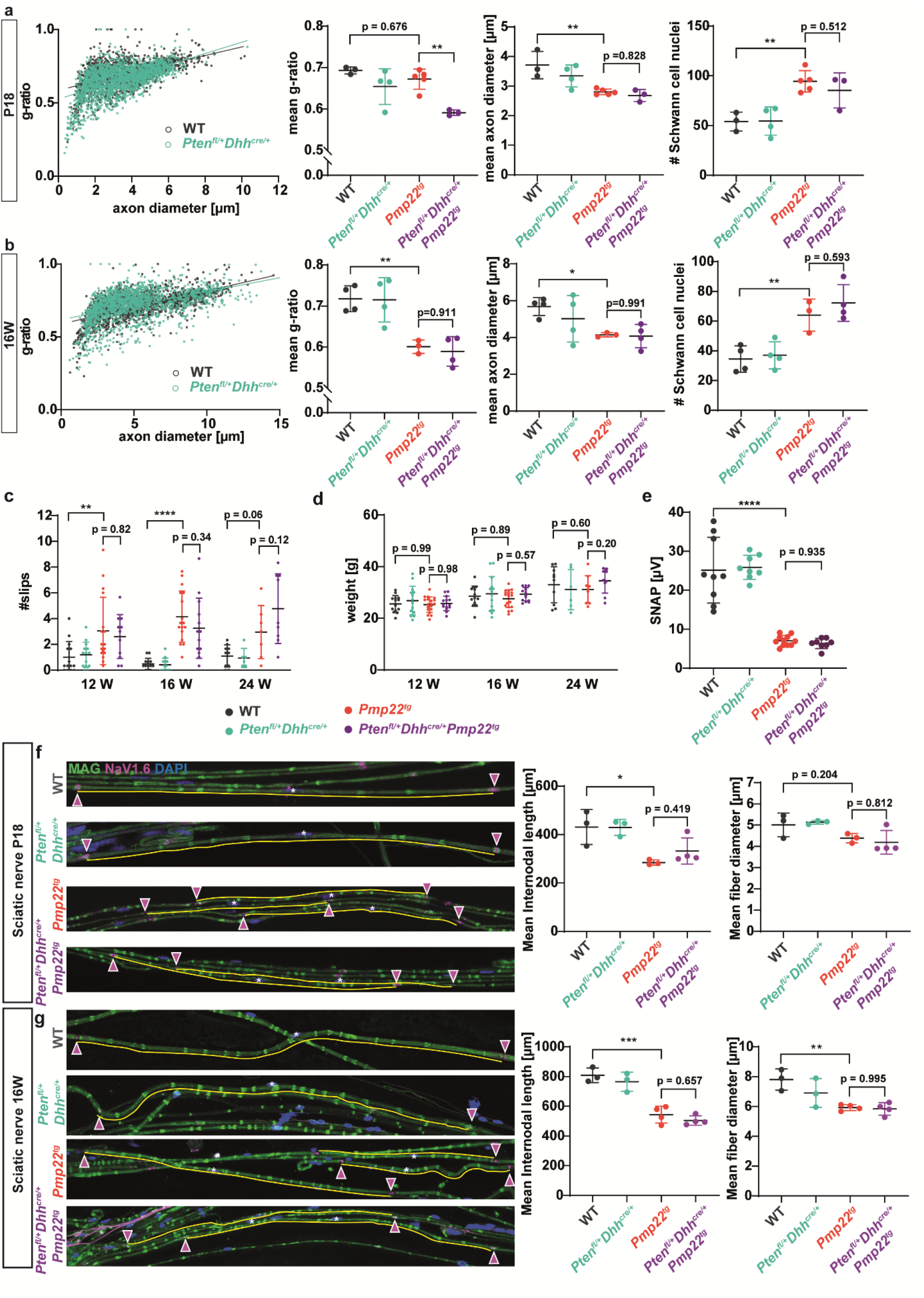
Unaltered internodal length and myelin sheath thickness in *Pten*^*fl/+*^*Dhh*^*cre/+*^ mice. **a** G-ratio plotted against axon diameter of wildtype (WT, grey) and *Pten*^*fl/+*^*Dhh*^*cre/+*^ (turquoise) femoral nerves at P18. Mean g-ratio is unaltered in *Pten*^*fl/+*^*Dhh*^*cre/+*^ and *Pmp22*^*tg*^ mice compared to WT controls and decreased in *Pten*^*fl/+*^*Dhh*^*cre/+*^*Pmp22*^*tg*^ mice (left panel). Mean axon diameters are reduced in *Pmp22*^*tg*^ and *Pten*^*fl/+*^*Dhh*^*cre/+*^*Pmp22*^*tg*^ mice (middle panel). The number of Schwann cell nuclei per femoral nerve cross section is increased in *Pmp22*^*tg*^ and *Pten*^*fl/+*^*Dhh*^*cre/+*^*Pmp22*^*tg*^ mice. WT n = 3, *Pten*^*fl/+*^*Dhh*^*cre/+*^ n = 4, *Pmp22*^*tg*^ n = 5 and *Pten*^*fl/+*^*Dhh*^*cre/+*^*Pmp22*^*tg*^ n = 3 animals. **b** G-ratio plotted against axon diameter of WT (grey) and *Pten*^*fl/+*^*Dhh*^*cre/+*^ (turquoise) femoral nerves at 16 weeks of age. Mean g-ratio is unaltered in *Pten*^*fl/+*^*Dhh*^*cre/+*^ mice compared to WT controls and decreased in *Pmp22*^*tg*^ and *Pten*^*fl/+*^*Dhh*^*cre/+*^*Pmp22*^*tg*^ mice (left panel). Mean axon diameters are reduced in *Pmp22*^*tg*^ and *Pten*^*fl/+*^*Dhh*^*cre/+*^*Pmp22*^*tg*^ mice (middle panel). The number of Schwann cell nuclei per femoral nerve cross section is increased in *Pmp22*^*tg*^ and *Pten*^*fl/+*^*Dhh*^*cre/+*^*Pmp22*^*tg*^ mice. WT n = 4, *Pten*^*fl/+*^*Dhh*^*cre/+*^ n = 4, *Pmp22*^*tg*^ n = 3 and *Pten*^*fl/+*^*Dhh*^*cre/+*^*Pmp22*^*tg*^ n = 4 animals. **c** The number of slips on the elevated beam is similarly increased in *Pmp22*^*tg*^ and *Pten*^*fl/+*^*Dhh*^*cre/+*^*Pmp22*^*tg*^ mice compared to wildtype controls at all time points. Behavioral analysis was done at 12, 16 and 24 weeks of age. WT n = 10-14, *Pten*^*fl/+*^*Dhh*^*cre/+*^ n = 6-14, *Pmp22*^*tg*^ n = 9-19 and *Pten*^*fl/+*^*Dhh*^*cre/+*^*Pmp22*^*tg*^ n = 9-14 mice were analyzed. **d** Neither the weight of *Pmp22*^*tg*^ nor *Pten*^*fl/+*^*Dhh*^*cre/+*^*Pmp22*^*tg*^ mice is altered compared to wildtype controls at 12, 16 and 24 weeks of age. **e** Sensory nerve action potential amplitudes (SNAP) are decreased in the tail of *Pmp22*^*tg*^ and *Pten*^*fl/+*^*Dhh*^*cre/+*^*Pmp22*^*tg*^ mice compared to wildtype controls. For electrophysiology measurements WT n = 10, *Pten*^*fl/+*^*Dhh*^*cre/+*^ n = 8, *Pmp22*^*tg*^ n = 11 and *Pten*^*fl/+*^*Dhh*^*cre/+*^*Pmp22*^*tg*^ n = 8 mice were analyzed. **f** Example images of teased fiber preparations of WT, *Pten*^*fl/+*^*Dhh*^*cre/+*^, *Pmp22*^*tg*^ and *Pten*^*fl/+*^*Dhh*^*cre/+*^*Pmp22*^*tg*^ double mutants stained for MAG (green), NaV1.6 (magenta) and DAPI (blue) at P18. Internodes between two nodes (magenta arrowheads) are underlined in yellow and respective Schwann cell nuclei are marked by white stars. Mean internodal length (left panel) is significantly reduced in *Pmp22*^*tg*^ teased fibers compared to wildtype controls at P18, whereas *Pten*^*fl/+*^*Dhh*^*cre/+*^*Pmp22*^*tg*^ mice do not differ in internodal length compared to *Pmp22*^*tg*^ mice. Mean fiber diameters are not significantly altered (right panel). Analysis was performed on 100 internodes of n = 3-4 animals per group. **g**. Example images of teased fiber preparations of WT, *Pten*^*fl/+*^*Dhh*^*cre/+*^, *Pmp22*^*tg*^ and *Pten*^*fl/+*^*Dhh*^*cre/+*^*Pmp22*^*tg*^ double mutants stained for MAG (green), NaV1.6 (magenta) and DAPI (blue) at 16 weeks of age. Internodes between two nodes (magenta arrowheads) are underlined in yellow and respective Schwann cell nuclei are marked by white stars. Mean internodal length (left panel) and fiber diameter (right panel) are significantly reduced in *Pmp22*^*tg*^ teased fibers compared to wildtype controls at 16 weeks of age, whereas *Pten*^*fl/+*^*Dhh*^*cre/+*^*Pmp22*^*tg*^ mice do not differ in internodal length compared to *Pmp22*^*tg*^ mice. Analysis was performed on 100 internodes of n = 3-4 animals per group. Means are displayed ± standard deviation. Statistical analysis was done using one-way ANOVA with Sidak’s multiple comparison test (*p ≤ 0.05, **p ≤ 0.01, ***p ≤ 0.001, ****p ≤ 0.0001).

**Supplementary figure S3:**
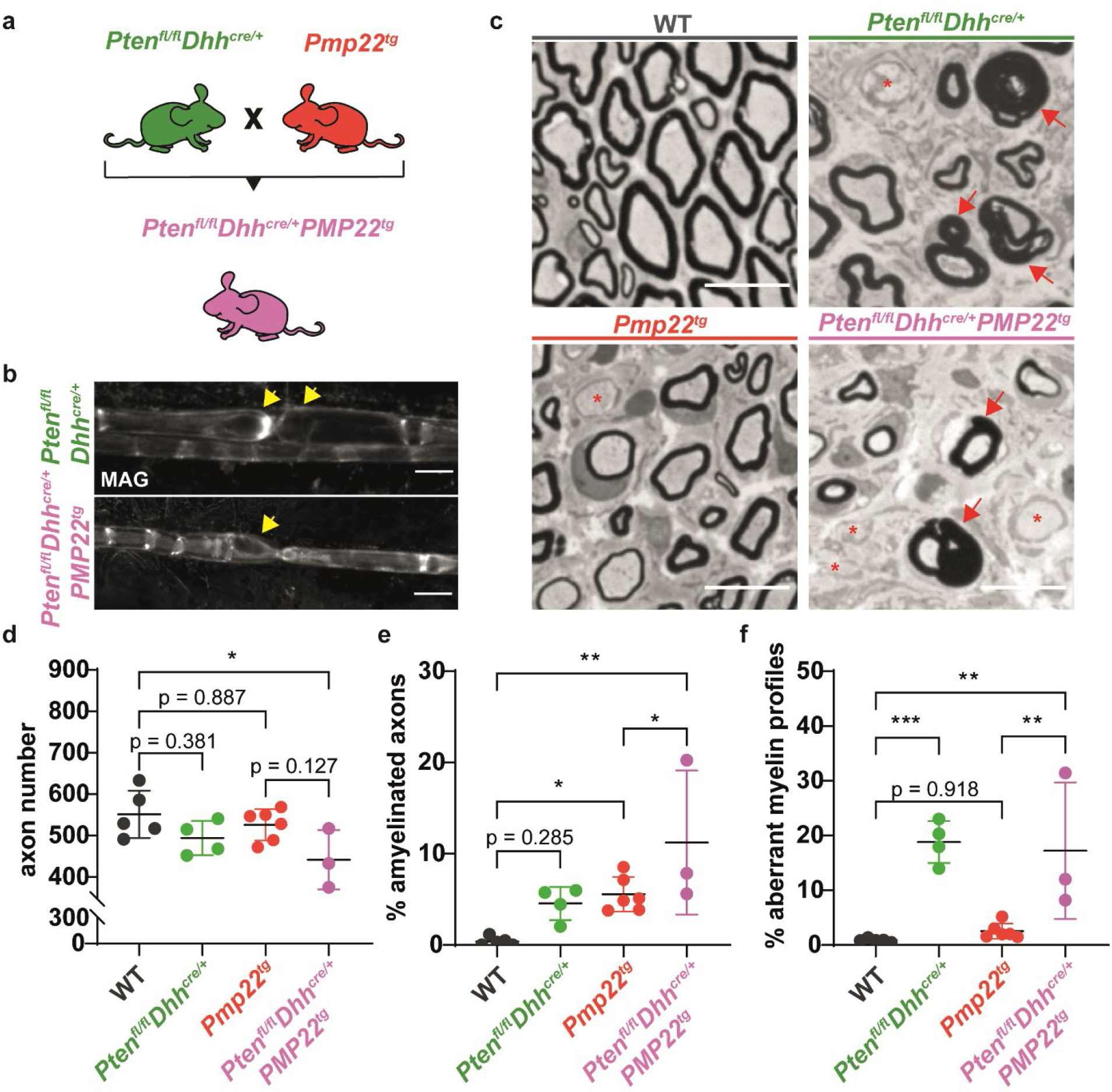
*Pten* depletion in *Pmp22*^*tg*^ mice leads to myelin abnormalities. **a** Crossing scheme of Schwann cell specific full Pten knockout mice (*PTEN*^*fl/fl*^*Dhh*^*cre/+*^) with CMT1A mice (*Pmp22*^*tg*^) to generate a full Pten knockout in CMT1A mice (*PTEN*^*fl/fl*^*Dhh*^*cre/+*^*Pmp22*^*tg*^). **b**. Teased fiber preparations of 8 weeks old *PTEN*^*fl/fl*^*Dhh*^*cre/+*^ (left panel) and *PTEN*^*fl/fl*^*Dhh*^*cre/+*^*Pmp22*^*tg*^ mice (right panel) show focal myelin thickening at paranodal loops as indicated by red arrows. Scale bar = 20 µm. **c** Semi-thin cross section of femoral nerves from WT, *PTEN*^*fl/fl*^*Dhh*^*cre/+*^, *Pmp22*^*tg*^ and *PTEN*^*fl/fl*^*Dhh*^*cre/+*^*Pmp22*^*tg*^ mice at 8 weeks of age. Red arrows indicate myelin abnormalities such as outfoldings and tomacula, asterisks indicate amyelinated axons. Scale bar = 10 µm. **d** Quantification of **c** displays reduced axon numbers in *PTEN*^*fl/fl*^*Dhh*^*cre/+*^*Pmp22*^*tg*^ mice compared to wildtype controls (left panel). The percentage of amyelinated axons is increased in *Pmp22*^*tg*^ and further elevated in *PTEN*^*fl/fl*^*Dhh*^*cre/+*^*Pmp22*^*tg*^ mice (middle panel). Pten depletion alone and in *Pmp22*^*tg*^ leads to an increase in axons with aberrant myelin profiles (right panel). WT n = 5, *PTEN*^*fl/fl*^*Dhh*^*cre/+*^ n = 4, *Pmp22*^*tg*^ n = 6 and *PTEN*^*fl/fl*^*Dhh*^*cre/+*^*Pmp22*^*tg*^ n = 3 animals were analyzed. Means are displayed ± standard deviation. Statistical analysis was done using one-way ANOVA with Sidak’s multiple comparison test (*p ≤ 0.05, ** p ≤ 0.01, ***p ≤ 0.001).

**Supplementary table 1:**
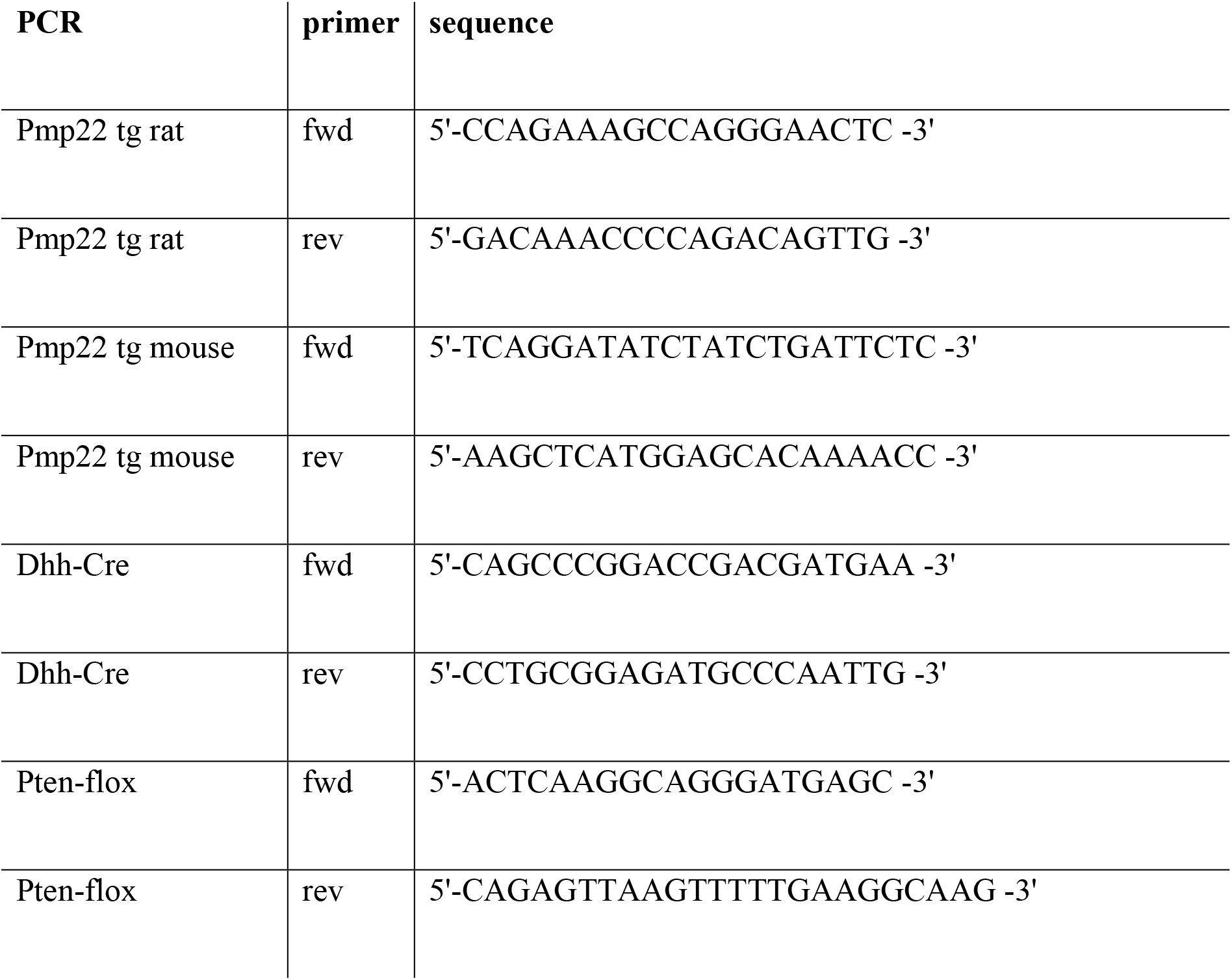
Genotyping primer.

**Supplementary table 2:**
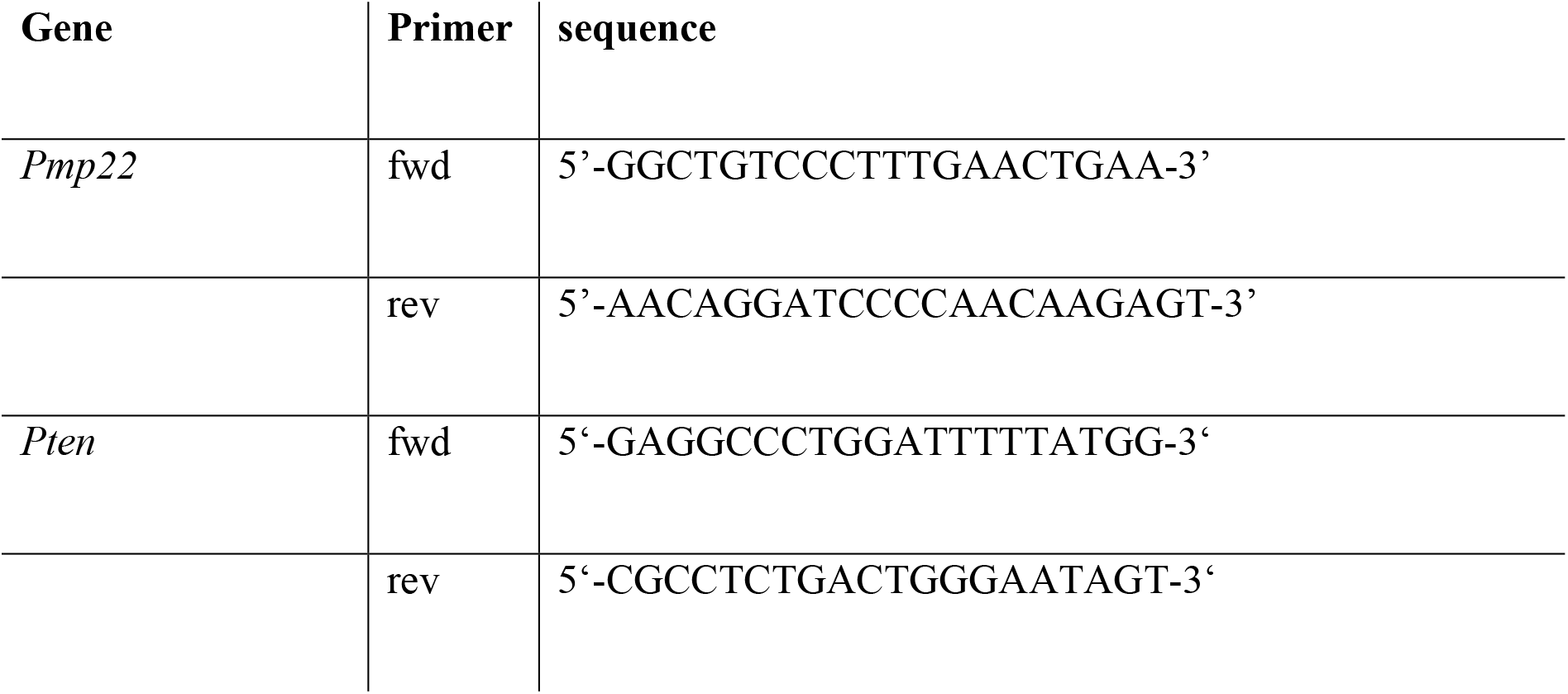
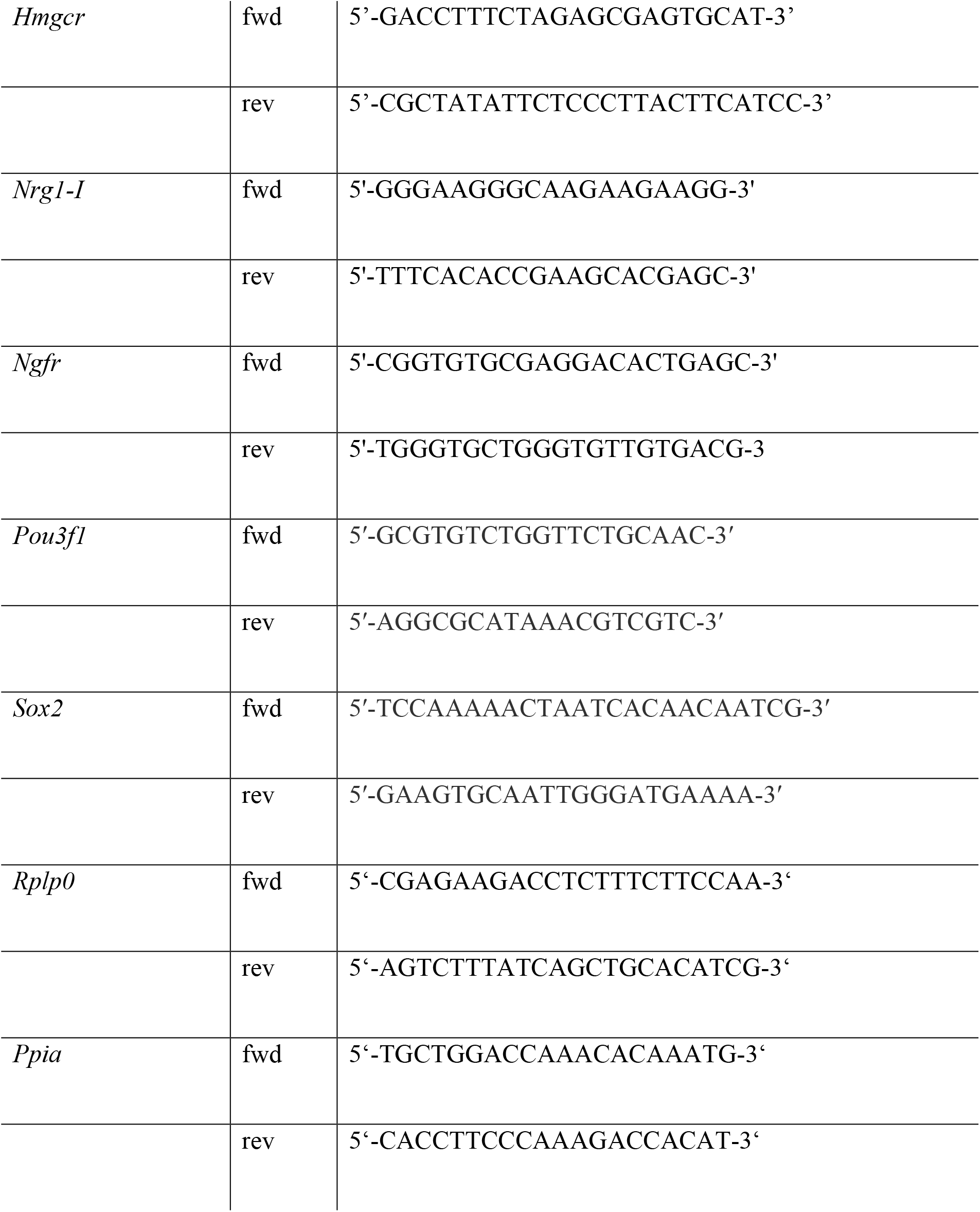
qRT-PCR primer.

## Notes

### Competing Interest Statement

The authors have declared no competing interest.

